# What Makes a Good Vaccine Antigen Target? Defining Key Features and Predicting Candidates in the *Staphylococcus aureus* Proteome

**DOI:** 10.64898/2026.08.09.743793

**Authors:** Nadia K. Prasetyo, Ries J. Langley, Fiona J. Radcliff, Paul P. Gardner

**Affiliations:** Department of Biochemistry, University of Otago, New Zealand; Faculty of Medical and Health Sciences, Department of Molecular Medicine & Pathology, University of Auckland, New Zealand

## Abstract

The rapid advancement of computational methods is transforming vaccine development by enabling faster, data-driven identification of promising antigens. In this study, we applied an in-silico pipeline to assess a broad set of sequence, structure, localisation, and immunology-derived features and determine which most effectively discriminate antigens from non-antigens in bacteria. Using these insights, we identified bacterial proteins with high potential as vaccine antigens. Applied to *Staphylococcus aureus*, this approach prioritized 304 candidate antigens, highlighting SSLs, nutrient acquisition factors, and cell wall-associated enzymes. While these findings demonstrate the potential of bioinformatics-guided antigen discovery, experimental validation remains essential. This work underscores the growing role of integrated computational and machine-learning approaches in accelerating next-generation vaccine design.

## Introduction

The rapid development and release of various SARS-CoV2 vaccines following the COVID-19 pandemic marked the beginning of a new era in vaccine development, characterized by advanced technology, accelerated design, and diverse vaccine platforms (1). Integrated computational approaches have since led to the identification of novel antigen targets for well-known pathogens, including *Mycoplasma genitalium*, oncogenic viruses, congenital Cytomegalovirus, and respiratory syncytial virus (2–5). Today, vaccines are being developed not only against viral pathogens but also against bacterial pathogens to combat rising antibiotic resistance, as cancer therapeutics, and for many other applications (6,7).

Successful vaccine design relies on the careful selection of appropriate antigens, whether derived from pathogens, synthetically produced, or structurally engineered, to induce strong antibody and T-cell responses (8). Advances in genomics and increasingly sophisticated machine learning (ML) tools have accelerated and refined vaccine target identification (1,9); however, significant challenges remain (10,11). Extensive pathogen diversity, the presence of multiple serotypes and strains, and rapid antigenic variation continue to hinder the development of vaccines capable of providing broad and long-lasting protection (8,12,13).

To address these challenges, immunoinformatics strategies such as reverse vaccinology and structural vaccinology have emerged as powerful tools for antigen discovery and vaccine design. Reverse vaccinology bypasses in-vitro pathogen culturing by identifying vaccine targets directly from genomic data (9,14). While structural vaccinology focuses on engineering or optimizing immunogens through detailed structural analysis and modeling of antigens. Recent advances in ML-based protein structure prediction and modeling have substantially accelerated both approaches (9,15). Nevertheless, ML-based methods remain highly dependent on the quality and quantity of available training data (9).

In this study, we employed an in silico approach to identify and evaluate bacterial proteins with potential as vaccine antigens. We developed a comprehensive bacterial antigen prediction pipeline that integrates biologically relevant feature analyses, including subcellular localisation, epitope prediction, conservation, allergenicity, immunogenicity, and both structural and compositional characteristics, utilizing a broad range of established bioinformatics tools. These features were statistically assessed across 12 representative bacterial pathogens for their ability to distinguish antigenic from non-antigenic proteins, forming the basis of a predictive model for antigen prioritization.

To demonstrate the practical application of this framework, we applied the resulting model to *Staphylococcus aureus* as a clinically relevant proof-of-concept pathogen. *S. aureus* is both a commensal and opportunistic bacterial pathogen that is a major cause of a wide range of clinical infections, including bacteremia and endocarditis (16,17). High rates of community- and hospital-acquired *S. aureus* infections, along with the widespread emergence of antibiotic resistance, underscores the urgent need for more effective therapeutic and preventative strategies (18–21). There are currently a number of *S. aureus* vaccines in clinical trials and pre-clinical stages, including live attenuated whole-cell vaccines, capsular polysaccharide targets, and multivalent protein subunit formulations (22,23). In addition, passive immunisation strategies using monoclonal antibodies have been investigated as an alternative method to manage *S. aureus* infections, with several candidates undergoing clinical trials (22,24). More recently, multivalent mRNA vaccines and novel recombinant protein subunit vaccines have shown promise in eliciting both humoral and cellular immune responses against *S. aureus* (25–27). Despite these extensive efforts, no licensed *S. aureus* vaccine is currently available for human use, highlighting the continued need to identify and evaluate novel vaccine antigens and strategies.

Within this context, we applied a random forest machine-learning model to rank the *S. aureus* proteome, resulting in the identification of 50 top-ranking proteins as promising vaccine antigen candidates.

## Methods

### Pathogen Sampling

Twelve bacteria were selected across bacterial phyla including bacillota, pseudomonadota, campylobacterota, spirochaetota, and chalmydiota (Figure 1). These organisms were chosen to provide broad taxonomic and high antigenic coverage, ensuring that the training set captures generalised features across major bacterial lineages. The selected species represent well-studied human pathogens for which substantial, high-quality antigen data exist in the Immune Epitope Database (IEDB) and the broader immunology literature. This emphasis on pathogens with established antigenic profiles helps ensure reliable annotation, reduces noise from poorly characterized organisms, and provides a strong foundation for downstream comparative analyses. Additionally, the set spans a wide range of clinically relevant bacteria, allowing the study to incorporate diverse host-pathogen interaction patterns (28). Selected bacteria includes *Streptococcus pneumoniae*, *Streptococcus pyogenes*, *Treponema pallidum*, *Chlamydia trachomatis*, *Neisseria gonorrhoeae*, *Helicobacter pylori*, *Brucella melitensis, Coxiella burnetii*, *Pseudomonas aeruginosa*, *Vibrio cholerae*, *Haemophilus influenzae*, and *Salmonella enterica* subsp. enterica serovar Enteritidis.

**Figure 1:**
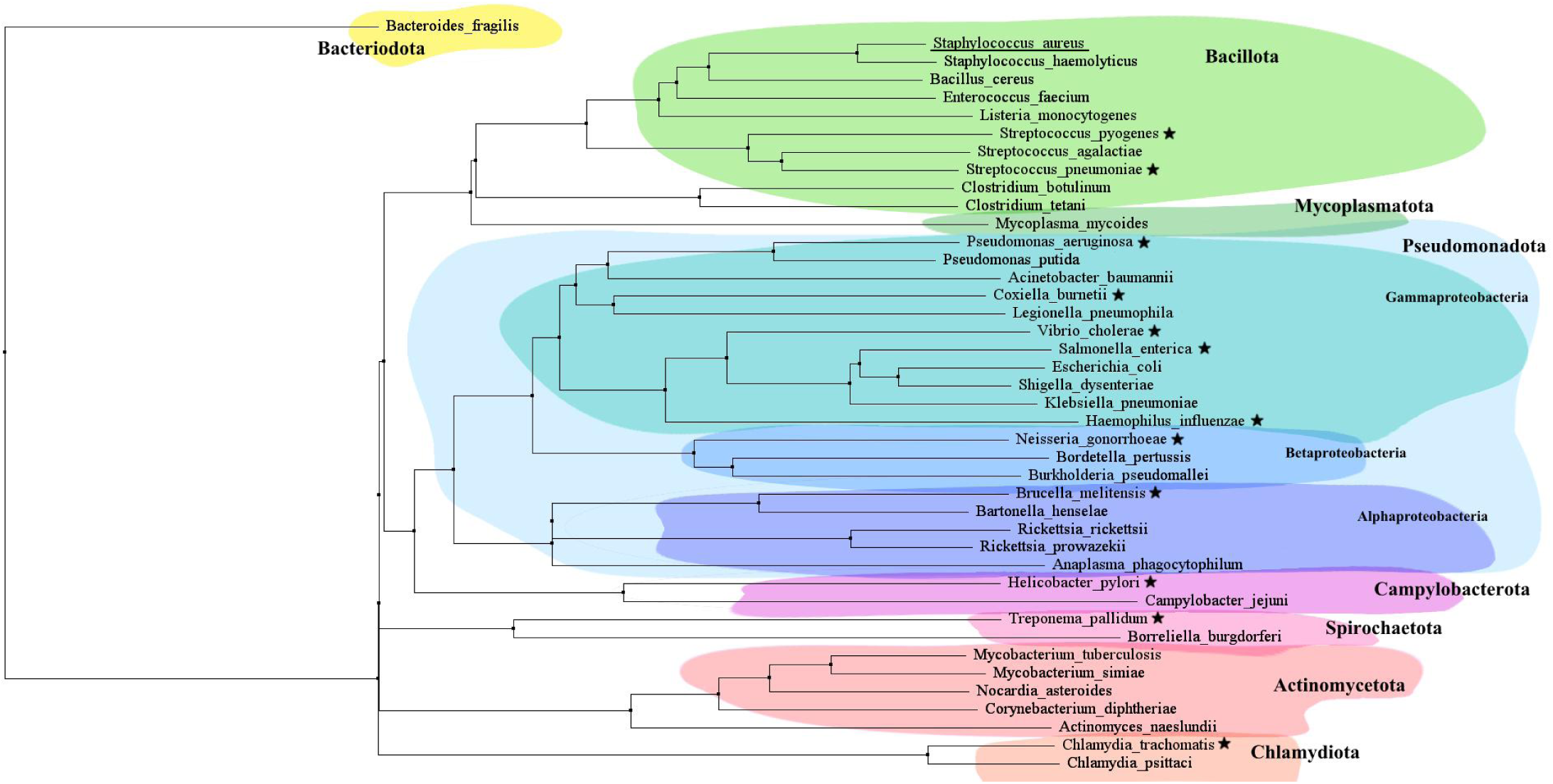
Phylogenetic tree of selected pathogenic bacteria. This is a representative phylogenetic tree of pathogenic bacteria and does not include all members of each phylum. The tree was constructed by performing a multiple sequence alignment of 16S rRNA sequences from the species shown. The 12 pathogens selected for feature evaluation and training are marked with a star (★). The target organism, *Staphylococcus aureus*, is underlined to highlight its position relative to the selected bacteria. Major bacterial phyla and classes are labeled and visually grouped using color shading for clarity.

The selection of these bacterial species was guided by several key considerations to ensure a balanced and biologically meaningful training set. First, the panel was designed to be broadly representative across the bacterial tree of pathogens rather than concentrated within a single lineage. This diversity reduces the risk of generating models that are overly tailored to a particular clade or its close relatives and minimizes bias that can arise from taxonomic over-representation. Second, avoiding over-representation of any single group reduces the potential for over-training and data-leakage, which is particularly important when antigenic features may be conserved along evolutionary lines (29,30). Third, accounting for phylogenetic relationships is crucial because evolutionary relatedness is a significant confounder in genomics and immunology research, leading to spurious correlations if not properly controlled (29,31). By incorporating phylogenetically diverse bacterial species, this study intentionally balances representation while preserving the evolutionary context necessary for robust, generalizable antigen prediction.

### Staphylococcus aureus Strain Selection

Six *S. aureus* strains were used to source the 6 Clonal Complexes that cover the majority of *S. aureus* infections: CC1, CC5, CC8, CC22, CC30, and CC93 (Supplementary Table 1). The genomes and proteomes of the strains were fetched from the NCBI genome datasets (32).

### Pathogen Genomes and Proteomes

Bacterial genomes and proteomes used in this study were derived from the [NCBI genome dataset](https://www.ncbi.nlm.nih.gov/datasets/genome/), downloaded via the [NCBI Datasets v2 REST API](https://www.ncbi.nlm.nih.gov/datasets/docs/v2/api/rest-api/) (32).

### Antigen Datasets

Antigens for each bacterium were sourced from the [IEDB](https://www.iedb.org/) (Last Updated: October 12, 2025) where experimental data on human B cell and T cell epitopes are cataloged. Additional antigens were directly sourced from published literature for each bacterium (Supplementary Table 2). Relevant scientific articles were compiled using the Litmaps search tool (version 2025-01-16), applying the keyword sets listed in Supplementary Table 3 (33). All searches were conducted between July and October 2025. Antigen protein and nucleotide sequences were fetched from the [UniProt website REST API](https://www.uniprot.org/api-documentation/uniprotkb) (Release 2025_04) (34). Subsequently, to get the antigen sequences for each strain, the antigen protein and nucleotide sequences were aligned to the bacterial proteomes and genomes using MMSeq2 (version 18.8cc5c) easy-search (35). The resulting bacterial strain-specific antigen sequences were used as the positive set for the concurrent analysis.

### Negative Dataset

A random selection of 200 proteins not previously known to be antigenic for a given bacterium was used as the negative set for the analysis. To control for potential confounding by sequence length, only proteins whose lengths fell within the range of the antigenic proteins for each bacterium were included. These non-antigen proteins are reviewed protein entries in [UniProtKB](https://www.uniprot.org/uniprotkb) (Release 2025_04) with the bacterium as the organism excluding the antigenic proteins listed as the positive dataset for the bacterium. To calculate protein and gene sequence conservation, 6 random complete genome assemblies were selected and downloaded from [NCBI genome dataset](https://www.ncbi.nlm.nih.gov/datasets/genome/) for a given bacterium. These presumed non-antigenic bacterial proteins provide a negative control for the feature analysis and subsequently antigen prediction. The positive to negative set ratio varies for each bacterium but averages to approximately 1: 2 across all the bacteria (Supplementary Table 4).

### Feature Selection

To capture a wide variety of features from the bacterial proteins, eighteen bioinformatic tools were utilised. A complete list of which can be found in Supplementary Table 6. The features used as fields in the antigen prediction matrix were categorized into the groups.

Subcellular localisation refers to the identification of membrane-bound and secreted proteins by predicting the localisation of each protein. Tools that were used to predict protein localisation are SignalP-5.0, TargetP-2.0, DeepTMHMM (version 1.2.1240), and DeepLocPro1.0.

Allergenicity is a feature that evaluates the potential allergenicity of the proteins by using AlgPred2. We use this to filter protein candidates with high allergenic potential to ensure safety.

Immunogenicity includes the assessment of the antigenicity of each protein using IFNepitope2 to prioritise proteins with interferon activation potential.

Conservation analysis involves the calculation of protein and DNA sequence conservation across multiple strains using MMseqs2 cluster, Rate4Site, and HyPhy dn/ds metrics such as FEL (Fixed Effects Likelihood), SLAC (Single-Likelihood Ancestor Counting), and FUBAR (Fast, Unconstrained Bayesian Approximation).

Epitope prediction includes tools that predict human T-cell and B-cell epitope peptides from the proteins. NetMHCPan4.2, NetMHCIIPan4.3, and MixMHC2pred for T-cell epitopes analyses the linear sequence of the protein to identify major histocompatibility complex (MHC) class I and class II epitopes. Seven representative MHCI corresponding human leukocyte antigen (HLA) alleles and seven MHCII corresponding HLA alleles (Supplementary Table 5) were used to maximise coverage.

BepiPred3.0 analyses the linear sequence of the proteins to predict B-cell epitopes while Ellipro and DiscoTope3.0 uses structural modeling to predict B-cell epitopes from the protein’s 3D structures. Epitope prediction tools help identify antigens based on the number of epitopes predicted, binding affinity, and coverage.

Finally, structure and composition analysis includes the identification and extraction of data from the protein structure and composition from the protein data bank (PDB) and AlphaFold Protein Structure Database V5. Tools that were used in structure and composition analysis includes ProtLearn, with a focus on amino acid indexes, and PyDSSP, a simplified implementation of the DSSP (Define Secondary Structure of Proteins) algorithm.

### Statistical Evaluation

To evaluate the significance of each feature, several statistical tests were run including Kolmogorov-Smirnov (KS) tests, receiver operating characteristic (ROC) analysis, standard directional Student’s t-tests, Spearman correlation, and a principle component analysis (PCA). Candidate features were filtered according to the KS difference statistic, p-value significance, and area under the ROC curve (AUROC). An overall PCA and Spearman correlation of the compilation of all bacterial data was then calculated and plotted to understand the importance of the features in classifying antigens from non-antigens.

### Staphylococcus aureus Protein Filtration

The *S. aureus* reference genome contains 2767 protein coding genes. This proteome was filtered to 1002 proteins based on likely subcellular localisation using SignalP, TargetP, and DeepLocPro. Proteins that were predicted to be localised in the cytoplasm and did not have any signal peptide or transmembrane domains were excluded according to the scores predicted by the tools. Subsequent feature analysis of the proteins were performed on the 6 *S. aureus* strains in Supplementary Table 1.

### Random Forest Algorithm

To estimate the likelihood of antigenicity, we developed a random forest algorithm using a python machine learning package scikit-learn (1.7.0) that evaluates 87 features for each protein accession, categorizing them into antigens and non-antigens. The model was trained using data from twelve diverse bacteria (from five phyla and seven classes, Figure 1) which includes 1,036 antigen annotated proteins and 2,018 likely non-antigen annotated length-matched proteins (we assume antigenicity is a relatively rare property), and evaluated on a smaller set of 183 antigen annotated proteins and 357 non-antigen annotated random proteins. The classification model demonstrated good performance on the evaluation dataset, achieving a validation AUC score of 0.92, indicating excellent precision and recall. The model was then used to rank 1002 *S. aureus* proteins, generating an antigen probability for each protein.

### Candidate Antigen Filtering Criteria

The candidate antigens predicted by the random forest model were further evaluated and filtered based on allergenicity, homology to human proteins, and predicted antigenicity. Allergenicity was assessed using AlgPred2, with the hybrid score serving as the filtering metric; proteins with a score above 0.3 were considered highly allergenic and excluded. Similarity to human proteins was evaluated using BLASTp against the human reference proteome (hg38, RefSeq: GCF_000001405.40). Proteins were removed if they showed >30% sequence identity with an e-value < 0.05, or if the BLASTp e-value was <10^-6^, ensuring minimal risk of cross-reactivity with human proteins (36). Finally, proteins sharing the same four-letter protein name abbreviation, or encoded by the same four-letter gene code, were grouped into homologous clusters and represented as a single entry. For each cluster, the mean predicted antigenicity and mean hybrid allergenicity score were calculated, and these aggregated values were used to determine the final ranking.

## Results

### Feature Importance Metrics

Among the 87 features captured from the bacterial proteins using eighteen bioinformatical software, some are better at distinguishing between the positive set and the negative set of proteins across the twelve bacterial species included. To statistically assess the contribution of each feature in distinguishing antigenic from non-antigenic proteins, we evaluated their distributions using the Kolmogorov-Smirnov (KS) test. Features with a KS test p-value below 0.05, indicating a significant difference between antigens and non-antigens, were ranked according to their KS difference statistic (Figure 2A). In addition, the discriminative performance of each feature was evaluated using the area under the receiver operating characteristic curve (AUROC). Only features with a KS p-value below 0.05 were included in figure 2B to ensure all AUROC values reflected performance above random expectation, the AUROC scores were adjusted using the formula max(AUROC, 1 - AUROC), setting the minimum possible value to 0.5. The directional bias of each feature was assessed using a two-sample t-test, and the results were indicated on the plots in figure 2 using hatching. A complete list of features, along with their corresponding KS and AUROC statistics, is provided in the Supplementary Material (Supplementary Figures 1 and 2, Supplementary Table 7). Additionally, to visualize and assess data variance, a principal component analysis (PCA) was performed on discriminating features (KS statistic > 0.15), and a biplot of PC1 and PC2 was generated (Supplementary Figure 3).

**Figure 2:**
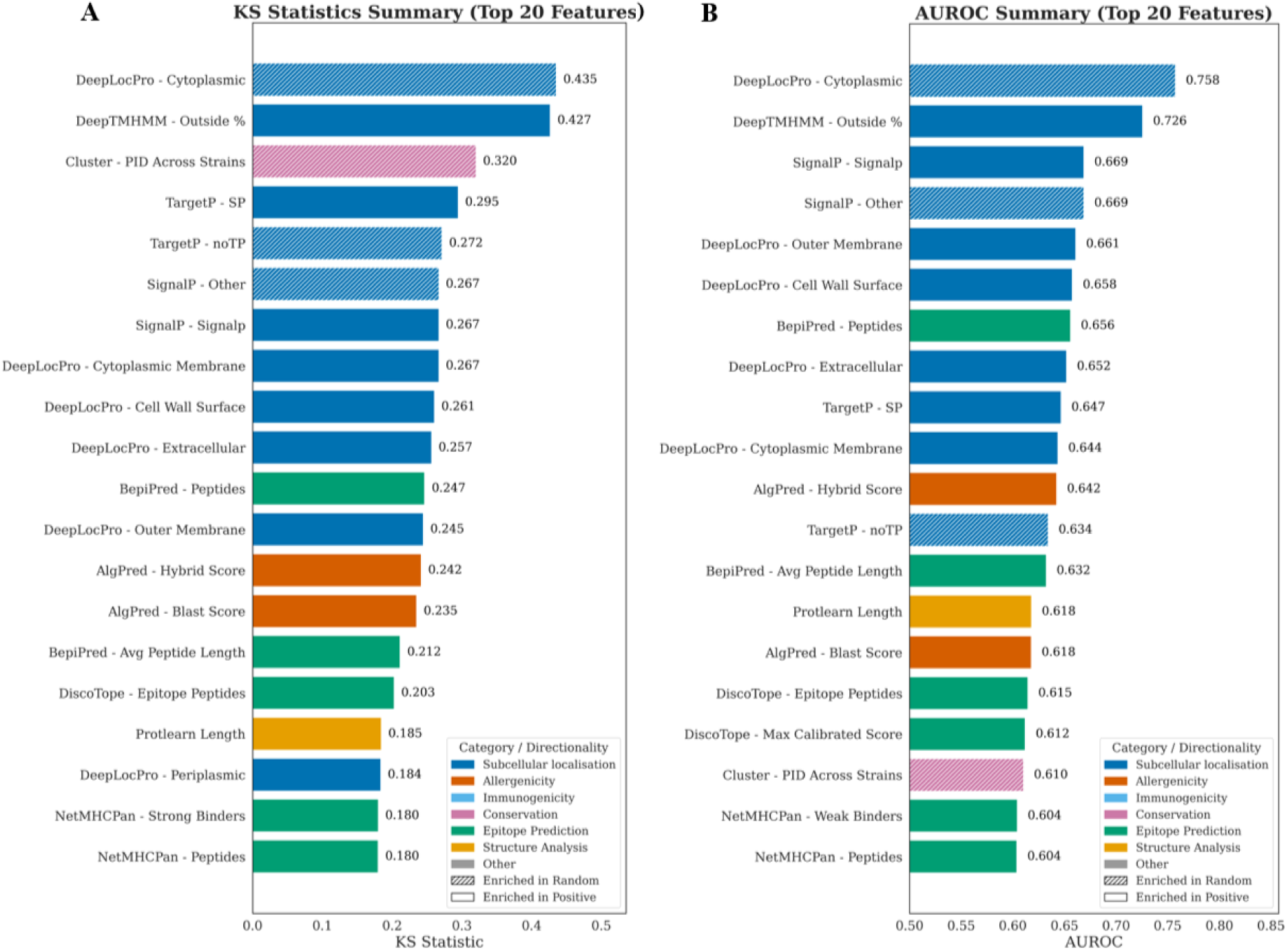
Overall Summary Statistics of the Top 20 Features (KS p-value < 0.05). **A)** KS statistic summary plot for the top 20 features according to the KS statistic value. **B)** AUROC summary plot for the top 20 features according to the adjusted AUROC value. Features are categorized into subcellular localisation (blue), allergenicity (vermillion), immunogenicity (sky blue), conservation (light pink), epitope prediction (green), structure analysis (yellow). Hatched patterned bars indicate enrichment of the feature in non-antigenic proteins (random set) while solid bars indicate enrichment in antigenic proteins (positive set).

We found that subcellular localisation features are the most informative predictors for antigenicity. The top 20 features were predominantly related to subcellular localisation, with features derived from DeepLocPro, DeepTMHMM, TargetP, and SignalP accounting for 11 of the top 20 features in the KS statistic summary and 10 of the top 20 in the AUROC summary (Figure 2). DeepLocPro cytoplasmic probability (prob_cytoplasmic) is shown to be the strongest discriminator between antigens and non-antigens, exhibiting a KS statistic of 0.435 (Figure 2A) and an AUROC of 0.758 (Figure 2B). This means that cytoplasmic localisation is negatively correlated with antigenicity. In addition to DeeplocPro cytoplasmic probability, several other features, cluster conservation score (sum of percent identity divided by the total number of strains), TargetP no signal peptide probability, and SignalP no signal peptide probability, were found to be consistently enriched in the non-antigenic bacterial proteins (random set) across all 12 bacterial species analyzed (Figure 2). All of these features thus are negatively correlated with antigenicity, indicating that if a protein were to be predicted to have cytoplasmic localisation, higher conservation between strains, or no signal peptides it is unlikely to be good antigen targets for vaccine development. On the other hand, antigens exhibited higher numbers of predicted B-cell epitopes, MHCI-binding peptides, and MHCII epitopes, along with higher DeepLocPro probabilities of outer membrane, cell wall surface, extracellular, cytoplasmic membrane localisation, and DeepTMHMM proportion outside. This suggests a positive correlation between antigenicity and extracellular localisation, membrane-associated proteins, cell-wall-surface localisation, the presence of a secretory peptide, enriched t-cell and b-cell peptides, and allergenicity. The enrichment of allergenicity in antigenic proteins may reflect molecular mimicry or immune-evasion strategies. This underscores the importance of carefully screening vaccine candidates for allergenic potential to minimize risks such as anaphylaxis or unintended autoimmune activation. At the same time, the enrichment of t-cell and b-cell epitope peptides in antigenic proteins suggests that these proteins possess more regions capable of interacting with host immune cells, thus triggering primary and secondary immune responses for successful vaccination response.

Interestingly, the KS and AUROC summary statistics also show that Protlearn length is included in the top 20 features in distinguishing between antigens and non-antigens. This suggests that there is a potential confounding factor regarding protein structure data availability, which may inflate the importance of structure-derived features such as protein structural data length (from ProtLearn). This is more likely a reflection of structural-data availability than a true biological signal. Antigens are more frequently represented in PDB or AlphaFold datasets, resulting in greater structural coverage and more derived structural descriptors overall.

The PCA shows no distinct clusters of proteins along either PC1 or PC2 (Supplementary Figure 3). However, there is a weak positive relationship between antigenicity and the PC1, as shown by more antigenic proteins (positive set) found towards higher values on the PC1 axis (Supplementary Figure 3). To understand this relationship, we investigated the loadings of the principal components 1 and 2. The loadings showed that PC1 is strongly influenced by epitope prediction features (mhci_num_peptides, mhcii_num_peptides, discotope), signal peptides (signalp_prob_signalp), and extracellular targeting features (targetp_prob_SP); in contrast, PC2 captured variation related to subcellular localisation and structural properties. This aligns with the findings of the KS test and AUROC tests, suggesting that antigens are more likely to have larger numbers of epitopes and signal peptides.

### Feature Correlations

To further understand the correlation of the different features, we investigated the Spearman correlation matrix and plotted it on a heatmap (Supplementary Figure 4). There are strong correlation clusters among features derived from the same prediction tools, reflecting redundancy among the different feature categories.

A large positively correlated cluster of features is formed from various structure-based features, including those derived from DiscoTope, ElliPro, DSSP, and ProtLearn (Figure 3). These features predict functionally distinct properties: subcellular localization, epitopes, and secondary structure data yet are unexpectedly positively correlated. Two potential confounding factors may contribute to this pattern. Firstly, these tools rely on the same class of input data: experimentally determined structures from the Protein Data Bank or predicted AlphaFold models, and therefore capture overlapping biophysical and surface-accessibility properties. This means that their reliance on the availability of PDB structural data is likely to affect the outputs of these tools. The bias is evident in the dataset itself: although the random protein set is approximately twice the size of the antigen set (2,397 vs. 1,239 proteins), both groups have nearly the same number of structural models (957 vs. 987) (Supplementary Table 11). Protein length may act as an additional confounder. The inclusion of ProtLearn’s length feature within this correlated cluster suggests that systematic differences in the length distributions of antigens and non-antigens could amplify correlations among structure-based features, particularly given that structural prediction quality and PDB coverage are themselves length-dependent. Together, these findings do not diminish the biological relevance of structural features, but highlight how structural tools can unintentionally encode data-driven biases that inflate feature correlations.

**Figure 3:**
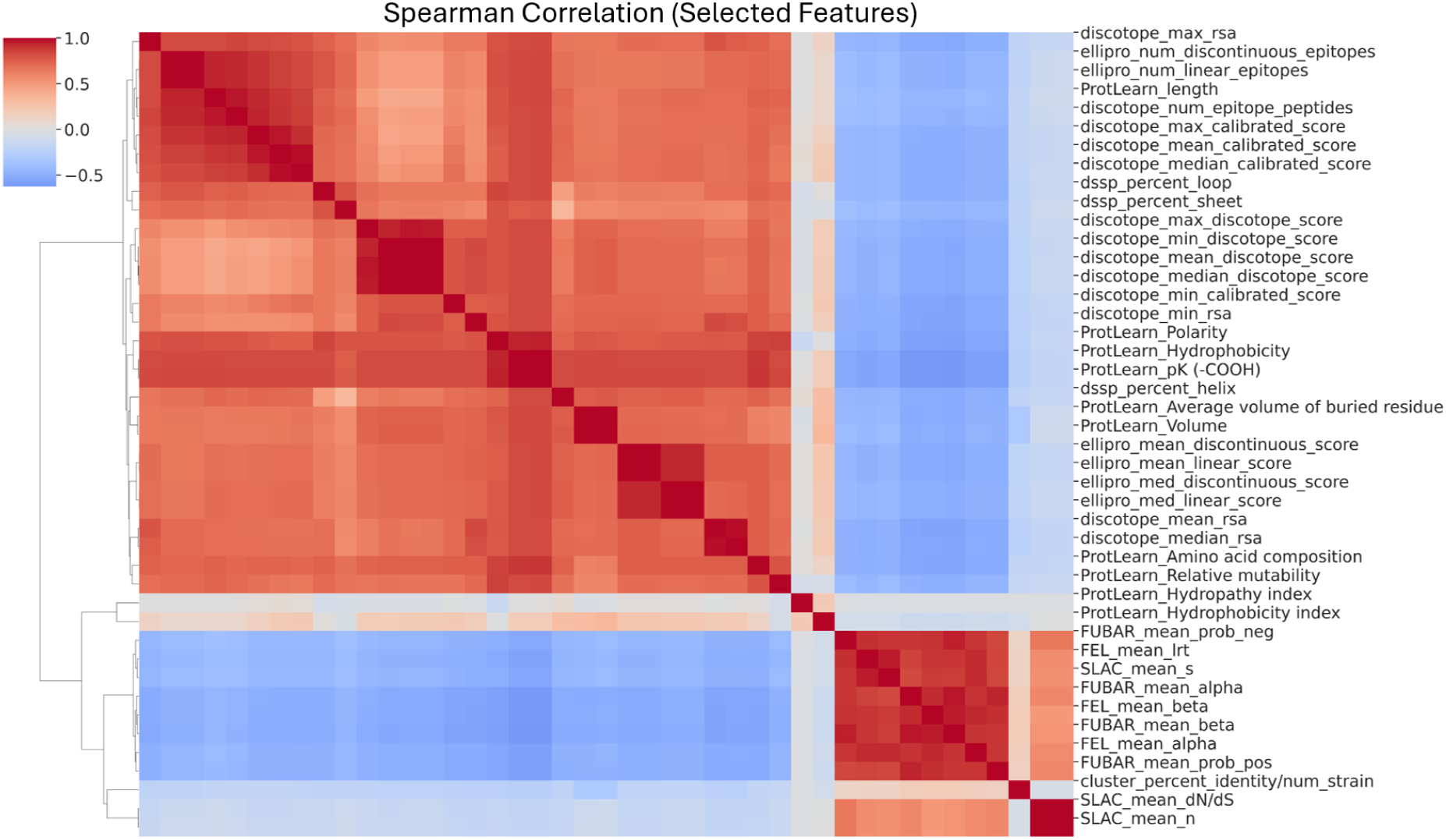
Spearman correlation matrix of selected protein features (clustered). A clustered heatmap of the Spearman rank correlation coefficients between all extracted protein features. The color scale represents the strength and direction of correlations, with red indicating positive correlations and blue indicating negative correlations. Hierarchical clustering was applied to both rows and columns to group features with similar correlation profiles.

A small cluster of positively correlated conservation features appears in the bottom right of figure 3, comprising dN/dS-based metrics (FEL, FUBAR, and SLAC) alongside a pairwise sequence alignment feature representing percent identity across antigen amino acid sequences. The dN/dS methods estimate evolutionary constraint at the nucleotide and amino acid level by comparing rates of synonymous and non-synonymous substitution, while percent identity provides a more direct measure of sequence similarity. Despite their methodological differences, the positive correlation among these features is expected: each captures a distinct aspect of evolutionary conservation, and their agreement suggests that conservation is a coherent, consistently detectable signal regardless of the metric used.

### Model Training and Prediction

Finally, we created a random forest classification model that was trained on antigenic and non-antigenic proteins from 12 pathogenic bacterial species and subsequently used to predict the most probable antigens within the *Staphylococcus aureus* proteome as an example case. The model was trained and evaluated using data that excluded *S. aureus* to ensure unbiased prediction performance. Predicted antigenic *S. aureus* proteins were further screened to minimize autoimmune risk by excluding those with high allergenicity (AlgPred hybrid score > 0.3) and those showing significant similarity to human proteins (37). Human-homologous proteins were identified using BLASTp with a threshold of >30% identity (e-value <0.05) or an e-value <1×10⁻⁶ (36). From the initial set of 1002 *S. aureus* proteins included in the random forest prediction, 210 antigens were removed due to high predicted allergenicity, 24 due to significant homology with human antigens, and 133 non-antigen predicted proteins (Supplementary Table 8-10). After filtering, antigens encoded by the same four-letter gene or protein code were grouped into homologous clusters and represented as single entries for downstream ranking and analysis, eliminating 374 redundant homologous antigens. This filtering and collapsing step trimmed down the list of predicted *S. aureus* antigens to 308 proteins, ranked by their predicted antigenicity. The top 100 filtered predicted antigenic *S. aureus* proteins are summarised and plotted in figure 4.

**Figure 4:**
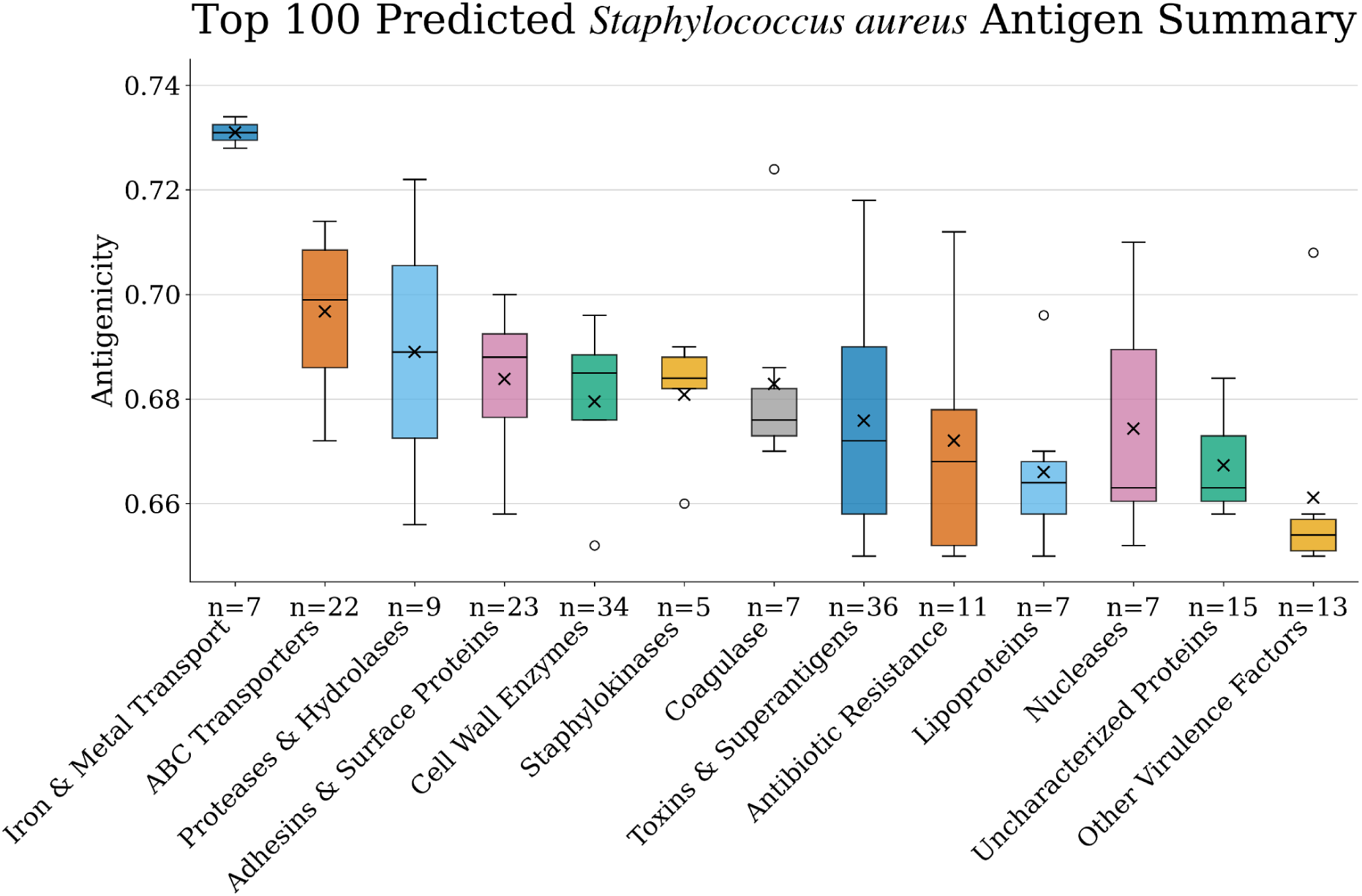
A summary plot of the top 100 predicted antigenic *S. aureus* proteins. The antigens included in the top 100 are categorised into 13 protein groups with the number of accessions (n) of each protein group labelled. Antigen probabilities were predicted using a random forest model trained on antigenic and non-antigenic protein data from 12 pathogenic bacterial species, excluding *S. aureus*.

Filtration of the predicted *S. aureus* antigens are critical, as they help ensure that any proposed vaccine antigens do not pose a risk of triggering severe allergic responses or autoimmune cross-reactivity, both major safety considerations in subunit vaccine development (8,38). The filtered antigens were therefore considered both safe and immunogenic. Of the 308 remaining proteins, the majority fell into eight broad functional categories, including toxins, immune evasion proteins, nutrient acquisition systems, cell wall and secreted enzymes, lipoproteins, and adhesins (Table 1). Together, these categories represent the multifaceted strategies by which *S. aureus* colonizes the host, acquires essential nutrients, avoids immune detection, and causes disease. Iron transporters such as IsdE and SirA returned the highest maximum antigenicity scores (0.734), followed by toxins including coagulase (0.724), and superantigen-like proteins (0.718). Notably, 96 proteins with predicted antigenicity scores up to 0.714 remained functionally uncharacterised. These findings reveal that *S. aureus* expresses a diverse array of surface-exposed and secreted proteins with high antigenic potential spanning critical virulence functions providing a rich pool of candidates for downstream experimental validation and multi-antigen vaccine design.

**Table 1:**
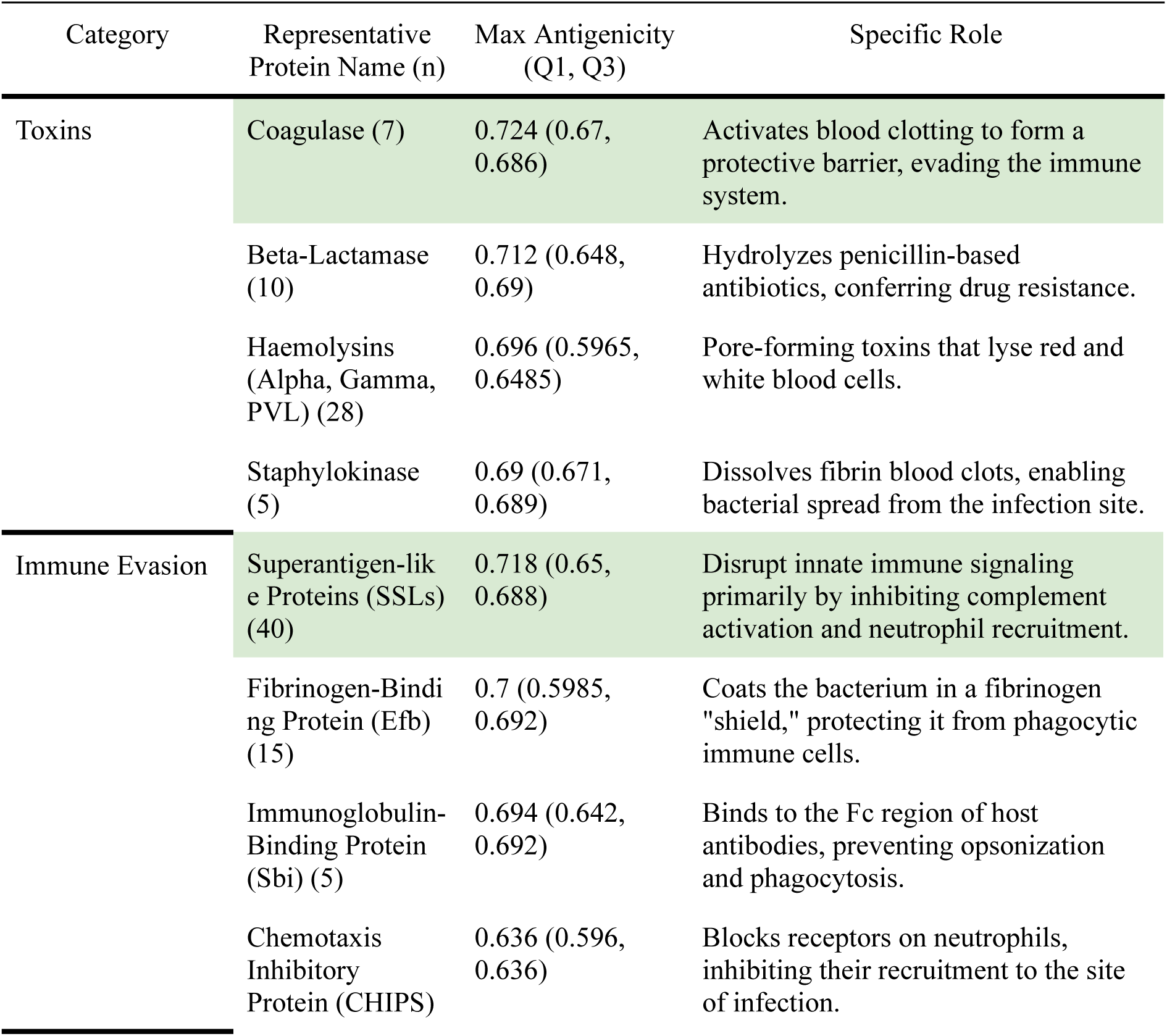
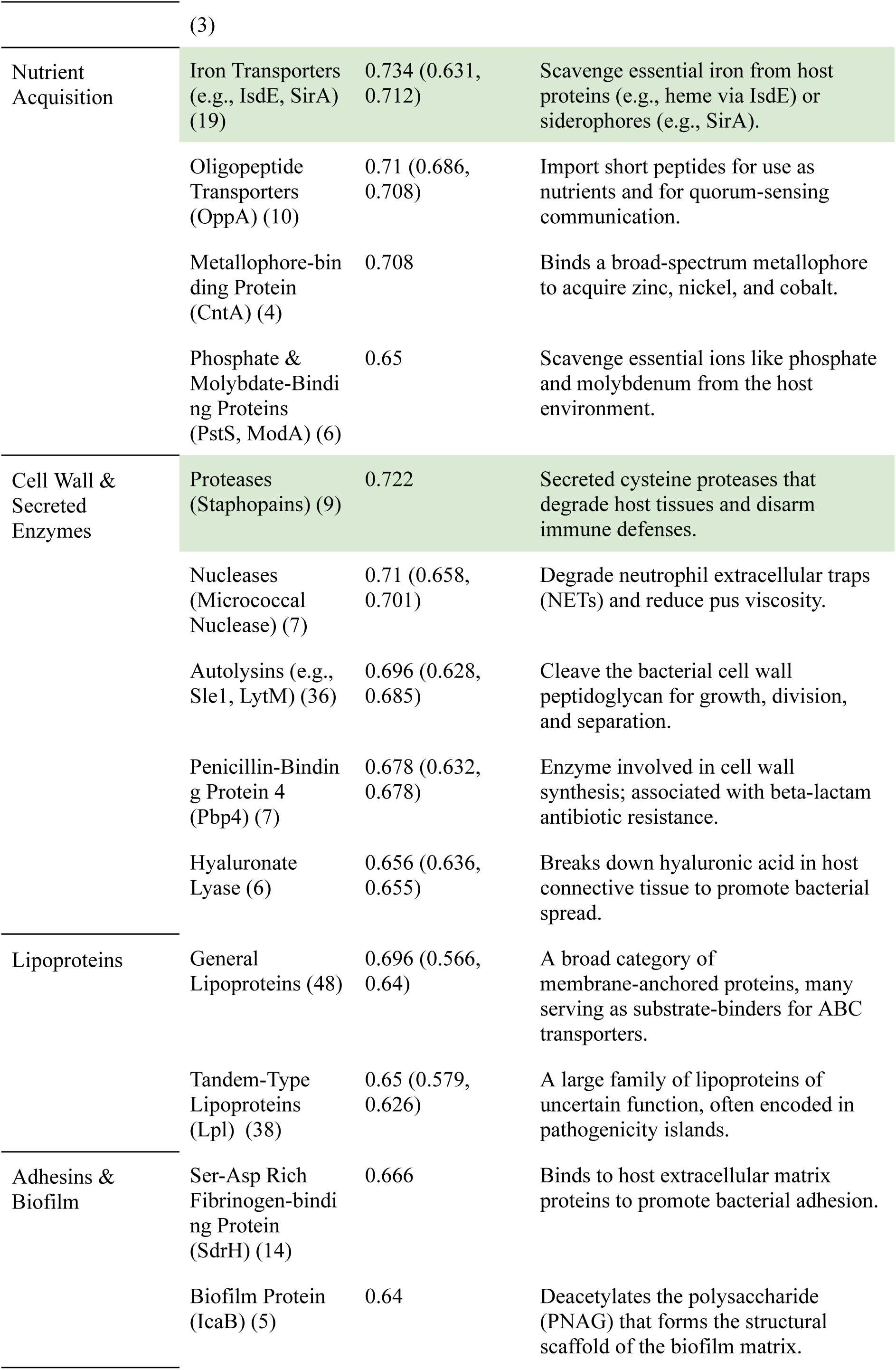
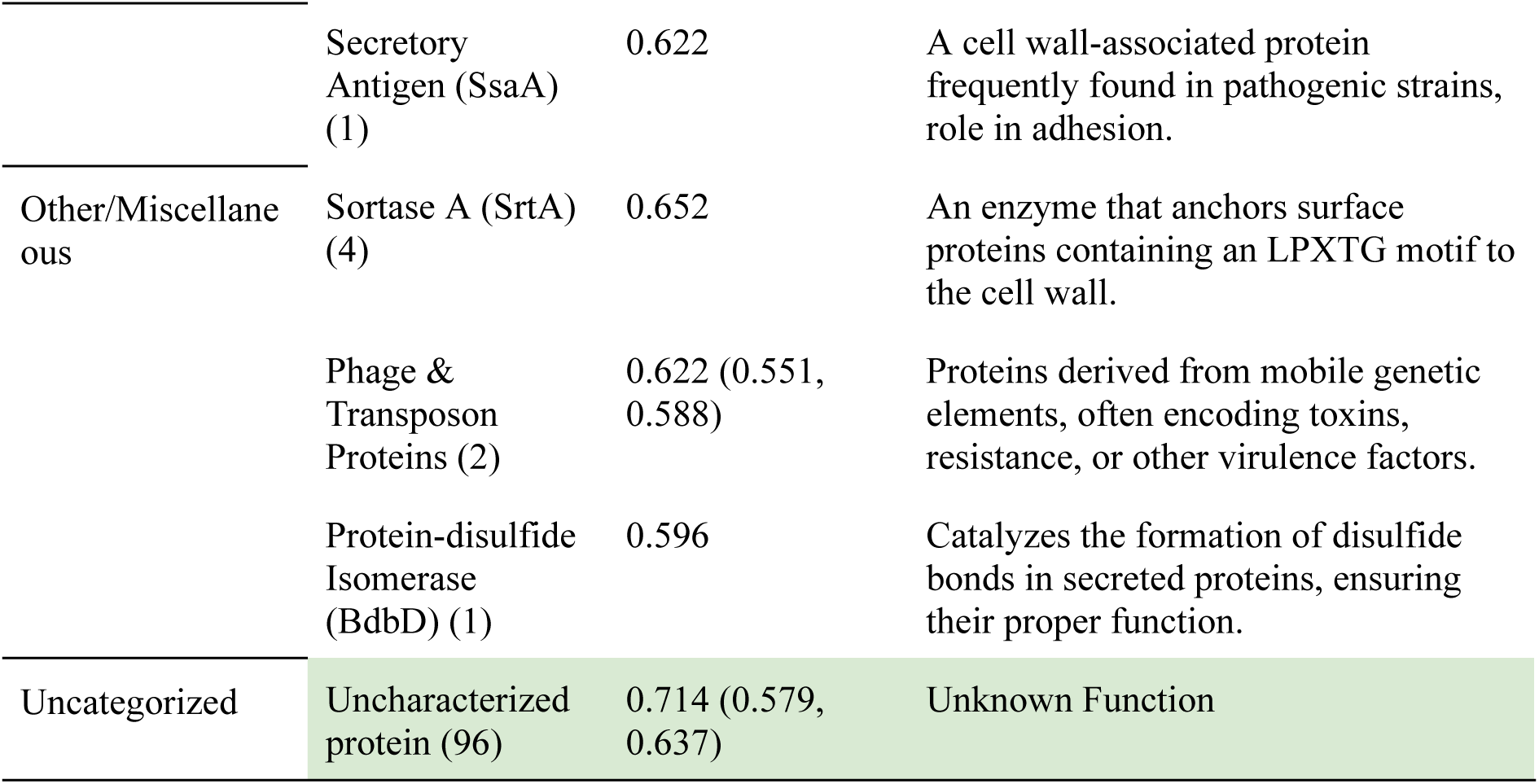
Summary categorization of the 308 predicted *S. aureus* antigen targets with a threshold predicted antigenicity of 0.5. The antigens are roughly categorized into 8 large categories labelled with representative names and number of accessions (n). The maximum predicted antigenicity, as well as lower (Q1) and upper quartile (Q3) were calculated for each representative protein group. The function and roles of each representative protein example in the categories are also included. Each category is sorted by max antigenicity. The top 5 max antigenicity representative proteins are highlighted in green.

### Essential Proteins

To further assess the biological relevance of the predicted antigens, all retained candidate proteins were cross-referenced with published *S. aureus* essential gene datasets and database of essential genes (Table 2) (39–44). This mapping step provides an additional layer of validation by identifying antigens that are not only surface-exposed and immunologically promising, but also required for bacterial viability. 8 proteins out of the predicted antigens are labeled as essential in *S. aureus*, including cell wall modification proteins (DltD), peptidoglycan remodeling proteins (IsaA, Sle1, PBP4), immune evasion proteins (Eap/Map), membrane biogenesis proteins (YidC), post-transcriptional regulatory proteins (RNase Y), and multifunctional proteins (Multifunctional Fusion Protein). Proteins encoded by essential genes are less likely to accumulate escape mutations and are typically conserved across strains, making them particularly attractive as broad-spectrum vaccine targets. However, essential proteins are often conserved across multiple pathogens and organisms, limiting their specificity as vaccine targets. For instance, protein DltD is part of the broadly conserved Dlt operon found across Gram-positive bacteria including *Lactobacillus* and other staphylococci (45). PBP4 belongs to a conserved class C low-molecular-weight PBP/DD-peptidase family found across diverse organisms including *Actinomadura* and *Escherichia coli* (46). YidC is conserved across all three domains of life, while RNase Y has orthologues in approximately 40% of sequenced eubacterial species (47,48). Although these four proteins are essential and predicted to be antigenic, they are less suitable as *S. aureus*-specific vaccine targets as immune responses generated against them may have off-target interactions, reducing the candidate antigen pool to 304 (49). This highlights the broader need for deeper investigation and experimental validation of predicted antigens before they can be considered viable vaccine candidates.

**Table 2:** Essential filtered predicted *S. aureus* antigens.

| Number of Accession | Maximum Antigen Probability | Protein Names |
| --- | --- | --- |
| 3 | 0.646 | Protein DltD |
| 5 | 0.646 | Immunodominant staphylococcal antigen A |
| 2 | 0.638 | N-acetylmuramoyl-L-alanine amidase sle1 |
| 4 | 0.622 | D-alanyl-D-alanine carboxypeptidase / Penicillin binding protein PBP4 |
| 3 | 0.622 | Multifunctional fusion protein |
| 1 | 0.622 | Extracellular adherence protein Eap/Map, MAP domain-containing protein |
| 2 | 0.54 | Foldase YidC |
| 1 | 0.516 | Ribonuclease Y |

## Discussion

Selecting appropriate vaccine antigens remains one of the central challenges in vaccine development. Effective vaccine antigens must be specific to the target pathogen, capable of eliciting protective immune responses, sufficiently safe to avoid triggering allergenic or autoimmune reactions, and ideally induce long-term immune memory (8,38). Here, we present a comprehensive in silico pipeline that addresses this challenge by statistically comparing antigen-associated features across twelve pathogenic bacteria, revealing clear biological patterns that distinguish antigenic from non-antigenic proteins without imposing prior assumptions about what defines an antigen.

Among the features examined, subcellular localisation emerged as the dominant computational signal distinguishing antigenic from non-antigenic proteins. This is consistent with the mechanistic basis of protective immunity, where surface-exposed and secreted proteins are are more accessible to innate and adaptive immune receptors, including pattern-recognition receptors, Toll-like receptors, T-cell receptors, and B-cell receptors and are therefore more likely to initiate and sustain an immune response (50). The co-enrichment of epitope prediction features in antigenic proteins reinforces this, reflecting the need to engage both humoral and cellular arms of the immune system for a durable vaccine response. Increased epitope diversity may further broaden immune recognition across hosts with diverse receptor repertoires, improving the likelihood of population-level protection (51).

The same surface exposure that makes a protein immunologically accessible, however, also subjects it to sustained host immune pressure, driving antigenic variation as a mechanism of immune evasion (52). This presents an inherent tension in antigen selection: while conserved proteins and epitopes are generally desirable for achieving broad strain coverage and limiting immune escape, the most accessible and immunogenic proteins are often those experiencing the strongest diversifying selection (53,54). This underlies the negative correlation between conservation and antigenicity observed in this study, and represents a fundamental challenge that must be considered when prioritising predicted antigens for vaccine development.

Although there are no currently licensed *Staphylococcus aureus* vaccines approved for human use, a number of *S. aureus* vaccines have reached clinical trials, including two that failed in clinical trials, and nine that are currently undergoing clinical trials in various stages (23,55). Although these candidate vaccines show robust antibody titres in vitro and early trials, they have consistently failed to translate to clinical protection in humans. The failures of previous clinical trials have been attributed to the complexity of *S. aureus*-host interactions, incomplete understanding of the immune mechanisms required for protection, and poor translation of animal model immunity to humans (23,55). Together, these challenges highlight the need for faster, more systematic approaches to antigen discovery that move beyond the conventionally targeted proteins.

The most frequently targeted antigens across ongoing and previous trials have been capsular polysaccharides CP5 and CP8, which featured in both the StaphVAX conjugate vaccine and Pfizer’s multivalent SA4Ag formulation, alongside surface-associated proteins including IsdB (Merck V710), ClfA, and MntC. More recent and ongoing trials have broadened the antigen repertoire to include toxins such as alpha-hemolysin (Hla) and Panton-Valentine leukocidin (PVL), as well as staphylococcal enterotoxin B (SEB) and protein A (SpA), reflecting a shift toward neutralising virulence factors alongside surface-targeting strategies (55). Encouragingly, several of these well-characterised antigens receive high predicted antigenicity scores in this study, including IsdB (0.734), ClfA (0.692), MntC (0.586), SpA (0.686), PVL (0.650), and Hla (0.666). This validates the model’s ability to correctly identify immunologically relevant proteins, as these surface-exposed or secreted proteins score highly on the same features (localisation, epitope density) that the model identifies as the strongest predictors of antigenicity. The clinical failure of previous trials with these antigens therefore highlights the complexity of translating immunogenicity into protective efficacy against a pathogen with sophisticated immune evasion strategies, rather than poor antigenicity (22,55). This is a limitation no computational model can resolve, and underscores the importance of complementing antigenicity prediction with rigorous experimental validation.

Beyond the antigens that have featured in clinical trials, other promising antigen candidates have been recently explored experimentally. A few notable examples include staphylococcal superantigen-like proteins (SSL3, SSL7, and SSL11) explored by Chen et. al, fibronectin binding protein A (FnBPA) fibronectin-binding protein A (FnBPA) in a multivalent mRNA-LNP formulation by Gao et. al, and a leukocidin AB (LukAB) vaccine formulation by Poolman et. al (26,27,56). Several of these are independently recovered by this pipeline: FnBPA (0.614), LukAB/LukGH (0.646), SSL3 (0.704) and SSL11 (0.664) rank prominently, providing independent computational support for their experimental prioritisation. Notably, SSL7 (0.678) and SEB (0.634) were excluded from the final ranked list due to predicted allergenicity, though their allergenic peptides may be engineered or optimized with rational design. In addition, the pipeline additionally identifies a large pool of high-scoring tandem-type lipoproteins and functionally uncharacterised proteins that would be unlikely to emerge from conventional antigen selection. These represent the class of candidates where a systematic computational approach adds the most value. Together with the validated known antigens, these candidates provide a rich and prioritised resource for downstream experimental investigation.

This study demonstrates the value of a comprehensive, computationally driven approach to antigen discovery. By integrating diverse biological features, including subcellular localisation, epitope prediction, conservation, allergenicity, and structural descriptors, we identified key properties that distinguish antigenic from non-antigenic bacterial proteins and applied these insights to prioritize 304 S. aureus vaccine targets.This work illustrates how computational pipelines can complement empirical approaches by recovering known immunogenic proteins while surfacing candidates that would be overlooked by conventional methods. A key limitation is that the training data encompasses a subset of human-associated bacterial pathogens, thus antigen prediction may be less reliable for organisms with poorly characterised virulence mechanisms or features not well represented in current databases. Future work should broaden the feature space and extend the framework to a wider range of pathogens, including viral species. Ultimately, this pipeline represents a principled and scalable first step in antigen discovery, not a replacement for experimental validation, which remains essential to confirm protective efficacy. As biological databases and predictive tools continue to improve, such frameworks will play an increasingly important role in accelerating vaccine development against both established and emerging infectious diseases.

## Data Availability

All complete genome annotated RefSeq genomes and proteomes used in this study are available for download from the NCBI Dataset (v15.0.0) (32,35). All protein sequences analysed, including antigens and non-antigens, are available in UniProtKB (Release 2025_04). Antigen names and genes reported in literature that were not found in the UniProtKB database were excluded from the analysis. The list of antigens included as positive sets in this study were sourced from both the IEDB and literature, and are available in the public GitHub repository (https://github.com/NadiaPrasetyo/mRNA_Target_Selection). PDB structures used in structural and composition analysis were sourced from the RCSB protein data bank (PDB) (https://www.rcsb.org/) through a text-based and sequence-based search using the RCSB PDB Search API (https://search.rcsb.org/index.html#search-api) (23,24). Proteins with no structures available in the RCSB PDB were recovered from the AlphaFold Protein Structure Database V5 (https://alphafold.com/) via the Alphafold UniProt accession based API (https://alphafold.com/#/). Essential genes for *Staphylococcus aureus* were compiled from multiple literature sources, including studies identifying genes required for growth in culture and during infection, as well as entries from the Database of Essential Genes (DEG) accessed through its web platform (http://www.essentialgene.org/) (39–44)

## Code Availability

The analysis of data, feature predictions, and *Staphylococcus aureus* antigen ranking prediction were done using conda 25.9.0, python 3.13.2, and R 4.4.3. The code used in the analysis is available from https://github.com/NadiaPrasetyo/mRNA_Target_Selection. The R packages used include: ggpattern 1.2.1, tidyverse 2.0.0, glue 1.8.0. Python packages used include: pandas 2.2.3, Requests 2.32.3, matplotlib 3.10.3, networkx 3.5, numpy 2.2.6, scipy 1.15.3, scikit-learn 1.7.0, tqdm 4.67.1, pybiolib 1.2.1022, torch 2.7.1+cpu, biopython 1.85, gemmi 0.7.3, openpyxl 3.1.5, pydssp 0.9.1, xgboost 3.0.4, mlxtend 0.23.4, protlearn 0.0.3, seaborn 0.13.2, adjustText 1.3.0, fair-esm 2.0.0. Additionally, various conda environments with defined packages and pip packages were used for various external bioinformatics tools. The Conda environments used throughout the analysis are detailed in the discotope_tools_dependencies.yml and ext_tools_dependencies.yml files in the GitHub repository. All of the external bioinformatics tools and software are available in their websites, GitHub, or as conda and pip packages. The complete list of external tools used in this study are: Conda packages including MMseqs2 (version 18.8cc5c), Rate4Site (version 2.01), [MAFFT](https://mafft.cbrc.jp/alignment/software/) (version 7.526), [HMMER](http://hmmer.org/) (version 3.4), [FastTree](http://www.microbesonline.org/fasttree/) (version 2.2.0), [MACSE](https://bioweb.supagro.inra.fr/macse/) (version 2.07), and [HyPhy](https://hyphy.org/) (version 2.5.83); Downloaded software including NetMHCPan4.2 (https://services.healthtech.dtu.dk/services/NetMHCpan-4.2/) NetMHCIIPan4.3 (https://services.healthtech.dtu.dk/services/NetMHCIIpan-4.3/), DeepLocPro1.0 (https://services.healthtech.dtu.dk/services/DeepLocPro-1.0/), TargetP-2.0 (https://services.healthtech.dtu.dk/services/TargetP-2.0/), SignalP-5.0 (https://services.healthtech.dtu.dk/services/SignalP-5.0/), BepiPred-3.0 (https://services.healthtech.dtu.dk/services/BepiPred-3.0/), MixMHC2pred2.0 (https://github.com/GfellerLab/MixMHC2pred), DiscoTope3.0 (https://github.com/mnielLab/discotope3_web), and IEDB ElliPro (http://tools.iedb.org/ellipro/); Downloaded database Pfam-A models (https://www.ebi.ac.uk/interpro/download/pfam/); Pip packages [DeepTMHMM](https://services.healthtech.dtu.dk/services/DeepTMHMM-1.0/) (via pybiolib version 1.2.1240), AlgPred2 (version 1.4), IFNepitope2 (version 1.2), [PyDSSP](https://github.com/ShintaroMinami/PyDSSP) (version 0.9.1), [ProtLearn](https://github.com/tadorfer/protlearn) (version 0.0.3), and [Gemmi](https://github.com/project-gemmi/gemmi) (version 0.7.3).

## Supplementary data

This work is accompanied by 11 supplementary tables and 4 supplementary figures, detailing the feature analysis, model performance, and full list of prioritized *S. aureus* antigen candidates.

## Acknowledgements

Nikki Moreland (Faculty of Medical and Health Sciences, University of Auckland), RNA & Data Science Platforms, Ministry of Business, Innovation and Employment, NZ.

