## Supplementary tables and figures. for "What Makes a Good Vaccine Antigen Target? Defining Key Features and Predicting Candidates in the *Staphylococcus aureus* Proteome"

### Supplementary Figures and Tables

**Supplementary Table 1: *S. aureus* strains selected for antigen prediction.**

| Strain | Accession/Reference number | NCBI RefSeq Genome Assembly | Clonal Complex |
|---|---|---|---|
| MSSA476 | BX571857 | GCF_000011525.1 | CC1 |
| N315 | BA000018 | GCF_000009645.1 | CC5 |
| NCTC 8325 | CP000253 | GCF_000013425.1 | CC8 |
| HO 5096 0412 | HE681097 | GCF_000284535.1 | CC22 |
| MRSA252 | BX571856 | GCF_000011505.1 | CC30 |
| JKD6159 | CP002114 | GCF_000144955.2 | CC93 |

**Supplementary Table 2: List of literature used to source additional antigens for each bacterium.**

| Bacteria | Additional Antigen Sources |
|---|---|
| *Staphylococcus aureus* | (1) |
| *Streptococcus pneumoniae* | (2–5) |
| *Streptococcus pyogenes* | (6–10) |
| *Pseudomonas aeruginosa* | (11–15) |
| *Chlamydia trachomatis* | (16–21) |
| *Neisseria gonorrhoeae* | (22–28) |
| *Coxiella burnetii* | (29–33) |
| *Helicobacter pylori* | (34–38) |
| *Brucella melitensis* | (39–42) |
| *Treponema pallidum* | (43–46) |
| *Vibrio cholerae* | (47–50) |
| *Haemophilus influenzae* | (51–53) |
| *Salmonella enterica* subsp. enterica serovar Enteritidis | (54–58) |

**Supplementary Table 3: Keywords used to browse scientific literature in Litmaps (Version 2025-01-16) [Search tool].**

| Keywords |
|---|
| *[bacterium]* virulence |
| *[bacterium]* antigen |
| *[bacterium]* vaccine |
| *[bacterium]* pathogenicity |

**Supplementary Table 4: Number of antigenic and non-antigenic sample proteins for the bacteria.**

| Bacteria | Number of proteins | |
|---|---|---|
| | Positive set | Negative Set |
| *Streptococcus pneumoniae* | 69 | 200 |
| *Streptococcus pyogenes* | 132 | 197* |
| *Pseudomonas aeruginosa* | 129 | 200 |
| *Chlamydia trachomatis* | 123 | 200 |
| *Neisseria gonorrhoeae* | 87 | 200 |
| *Coxiella burnetii* | 148 | 200 |
| *Helicobacter pylori* | 32 | 200 |
| *Brucella melitensis* | 138 | 200 |
| *Treponema pallidum* | 136 | 200 |
| *Vibrio cholerae* | 140 | 200 |
| *Haemophilus influenzae* | 77 | 200 |
| *Salmonella enterica* subsp. enterica serovar Enteritidis | 28 | 200 |
| Total | 1239 | 2397 |

* 3 of the 200 selected proteins failed to fetch from the UniProt and were excluded

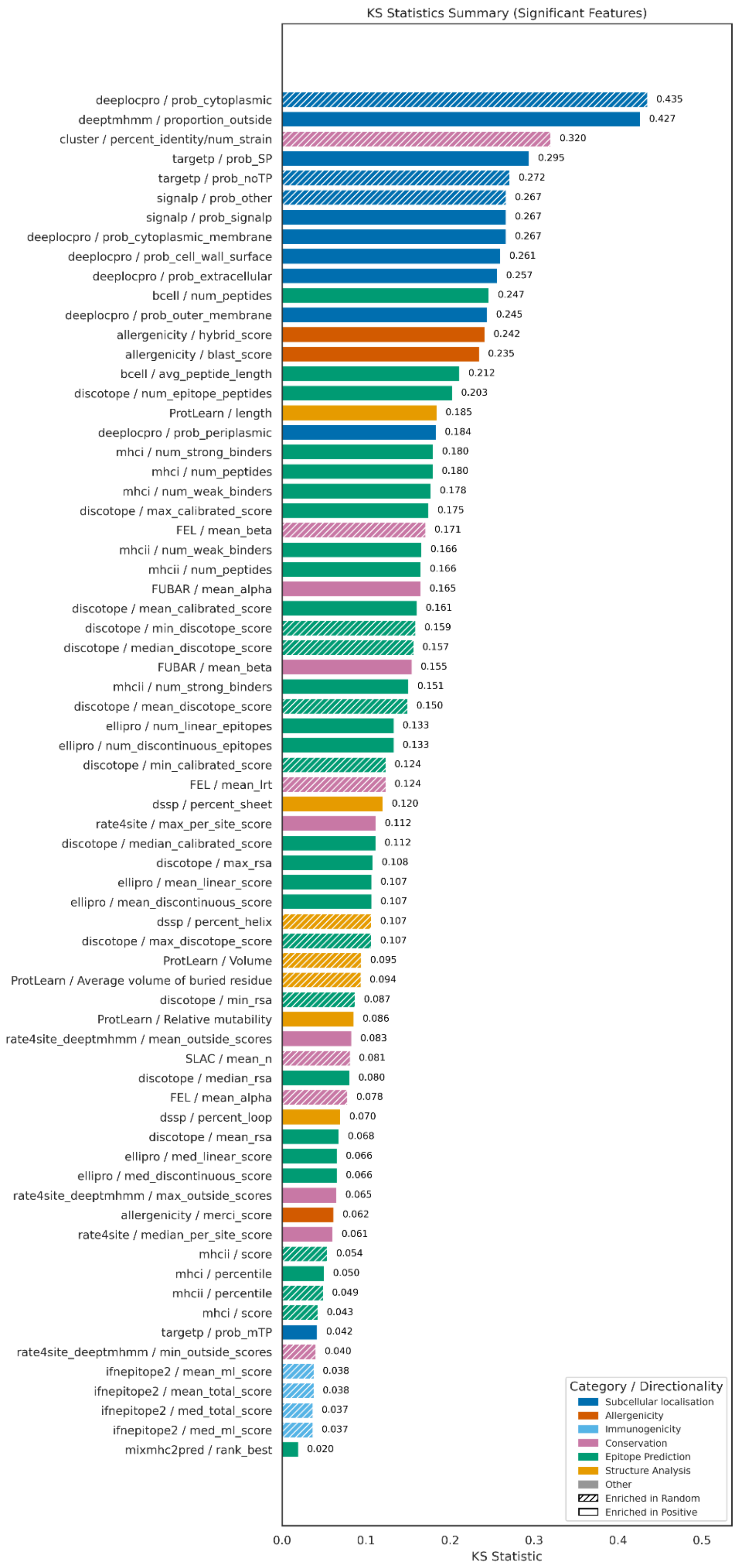


**Supplementary Figure 1: Complete KS statistical summary plot for all features (p-value < 0.5).**

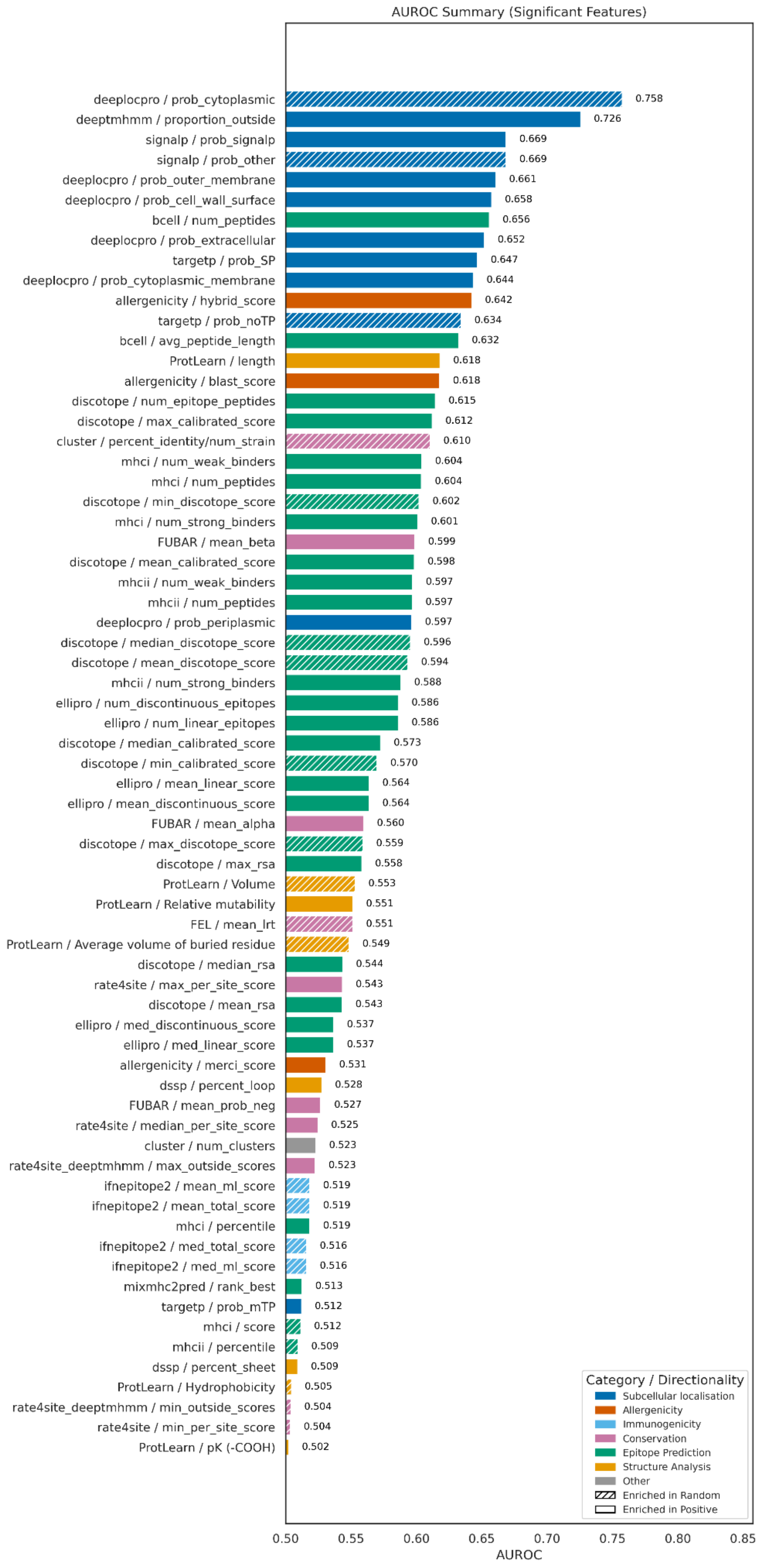


**Supplementary Figure 2: Complete AUROC statistical summary plot for all features (p-value < 0.5).**

**Supplementary Table 5. Human leukocyte antigen (HLA) alleles used in this study. MHC class I alleles were analysed using NetMHCpan 4.2, and MHC class II alleles were analysed using NetMHCIIpan 4.3 and MixMHCpred 2.**

| MHC Class | Alleles |
|---|---|
| **Class I** | HLA-A01:01, HLA-A02:01, HLA-A02:03, HLA-A02:06, HLA-A03:01, HLA-A11:01, HLA-A23:01, HLA-A24:02, HLA-A26:01, HLA-A30:01, HLA-A30:02, HLA-A31:01, HLA-A32:01, HLA-A33:01, HLA-A68:01, HLA-A68:02, HLA-B07:02, HLA-B08:01, HLA-B15:01, HLA-B35:01, HLA-B40:01, HLA-B44:02, HLA-B44:03, HLA-B51:01, HLA-B53:01, HLA-B57:01, HLA-B58:01 |
| **Class II** | HLA-DRB1:0301, HLA-DRB1:0701, HLA-DRB1:1501, HLA-DRB3:0101, HLA-DRB3:0202, HLA-DRB4:0101, HLA-DRB5:0101 |

**Supplementary Table 6: Complete list of features and tools used in this study.**

| Category | Feature | Bioinformatical Tool | Referen ce |
|---|---|---|---|
| Allergenicity | Predicted protein allergenicity (hybrid score) | AlgPred2.0 | (59) |
| | Predicted protein allergenicity based on BLAST similarity | AlgPred2.0 | |
| | Predicted allergenicity from MERCI patterns | AlgPred2.0 | |
| Conservation | Conservation across strains (percent identity / number of strains) | Mmseqs2 | (60) |
| | Number of sequence clusters across strains | MMseqs2 | |
| | dN/dS selection pressure estimate (SLAC mean s) | SLAC | (61,62) |
| | dN/dS selection pressure estimate (SLAC dN/dS) | SLAC | |
| | dN/dS selection pressure estimate (SLAC mean n) | SLAC | |
| | dN/dS FEL likelihood ratio test (selection estimation) | FEL | |
| | dN/dS FEL beta parameter (selection estimate) | FEL | |
| | dN/dS FEL alpha parameter (selection estimate) | FEL | |
| | dN/dS FUBAR posterior probability of positive selection | FUBAR | (61,63) |
| | dN/dS FUBAR beta parameter (selection estimate) | FUBAR | |
| | dN/dS FUBAR alpha parameter (selection estimate) | FUBAR | |
| | dN/dS FUBAR posterior probability of negative selection | FUBAR | |
| | Rate4Site conservation score in extracellular regions (mean) | Rate4Site + DeepTMHMM1.0 | (64,65) |
| | Rate4Site conservation score in extracellular regions (maximum) | Rate4Site + DeepTMHMM1.0 | |
| | Rate4Site conservation score in extracellular regions (minimum) | Rate4Site + DeepTMHMM1.0 | |

| | | | |
|---|---|---|---|
| | Rate4Site maximum site conservation score | Rate4Site | (64) |
| | Rate4Site minimum site conservation score | Rate4Site | |
| | Rate4Site median site conservation score | Rate4Site | |
| | Rate4Site mean site conservation score | Rate4Site | |
| | Rate4Site positions lacking score (NaN count) | Rate4Site | |
| Epitope prediction | Number of predicted B-cell epitope peptides | BepiPred3.0 | (66) |
| | Average length of predicted B-cell epitope peptides | BepiPred3.0 | |
| | Mean surface accessibility (Discotope RSA) | DiscoTope3.0 | (66) |
| | Median surface accessibility (Discotope RSA) | DiscoTope3.0 | |
| | Minimum surface accessibility (Discotope RSA) | DiscoTope3.0 | |
| | Maximum surface accessibility (Discotope RSA) | DiscoTope3.0 | |
| | Number of predicted epitope regions (Discotope) | DiscoTope3.0 | |
| | Minimum calibrated epitope score (Discotope) | DiscoTope3.0 | |
| | Maximum calibrated epitope score (Discotope) | DiscoTope3.0 | |
| | Mean calibrated epitope score (Discotope) | DiscoTope3.0 | |
| | Median calibrated epitope score (Discotope) | DiscoTope3.0 | |
| | Maximum epitope (Discotope) score | DiscoTope3.0 | |
| | Minimum epitope (Discotope) score | DiscoTope3.0 | |
| | Mean epitope (Discotope) score | DiscoTope3.0 | |
| | Mean discontinuous epitope score (ElliPro) | ElliPro | (67) |
| | Mean linear epitope score (ElliPro) | ElliPro | |
| | Median discontinuous epitope score (ElliPro) | ElliPro | |
| | Median linear epitope score (ElliPro) | ElliPro | |
| | Number of predicted linear B-cell epitopes (ElliPro) | ElliPro | |

| | | | |
|---|---|---|---|
| | Number of predicted discontinuous epitopes (ElliPro) | ElliPro | |
| | Number of predicted MHCI weak binders | NetMHCPan4.2 | (68) |
| | Number of predicted MHCI peptides | NetMHCPan4.2 | |
| | Number of predicted strong MHCI binders | NetMHCPan4.2 | |
| | MHCI binding percentile score | NetMHCPan4.2 | |
| | MHCI overall binding score | NetMHCPan4.2 | |
| | Number of predicted MHCII weak binders | NetMHCIIPan4.3 | (69) |
| | Number of predicted MHCII peptides | NetMHCIIPan4.3 | |
| | Number of predicted strong MHCII binders | NetMHCIIPan4.3 | |
| | MHCII binding percentile score | NetMHCIIPan4.3 | |
| | MHCII overall binding score | NetMHCIIPan4.3 | |
| Immunogenicity | Predicted IFN-inducing epitope score (mean total) | IFNepitope2 | (70) |
| | Predicted IFN-inducing epitope score (mean ML) | IFNepitope2 | |
| | Predicted IFN-inducing epitope score (median ML) | IFNepitope2 | |
| | Predicted IFN-inducing epitope score (median total) | IFNepitope2 | |
| | IFN epitope BLAST similarity score (mean) | IFNepitope2 | |
| | IFN epitope BLAST similarity score (median) | IFNepitope2 | |
| Structure & Composition | Protein length | ProtLearn | (71) |
| | Average buried residue volume | ProtLearn | |
| | Protein volume | ProtLearn | |
| | Relative mutability index | ProtLearn | |
| | Amino acid composition profile | ProtLearn | |
| | Hydrophobicity index | ProtLearn | |
| | Polarity index | ProtLearn | |
| | Hydrophobicity value | ProtLearn | |

| | | | |
|---|---|---|---|
| | Acidic pK (carboxyl group) | ProtLearn | |
| | Hydropathy index | ProtLearn | |
| | General sequence composition | ProtLearn | |
| | Average flexibility index | ProtLearn | |
| | Percent helix (secondary structure) | pyDSSP | (72,73) |
| | Percent sheet (secondary structure) | pyDSSP | |
| | Percent loop (secondary structure) | pyDSSP | |
| Subcellular Localisation | Probability of cytoplasmic localisation | DeepLocPro1.0 | (74) |
| | Probability of extracellular secretion | DeepLocPro1.0 | |
| | Probability of cytoplasmic membrane localisation | DeepLocPro1.0 | |
| | Probability of outer membrane localisation | DeepLocPro1.0 | |
| | Probability of cell wall surface localisation | DeepLocPro1.0 | |
| | Probability of periplasmic localisation | DeepLocPro1.0 | |
| | Proportion of residues predicted outside membrane | DeepTMHMM1.0 | (65) |
| | Probability of signal peptide (SignalP) | SignalP5.0 | (75) |
| | Other signal peptide prediction score | SignalP5.0 | |
| | Predicted secretory (SP) targeting peptide | TargetP2.0 | (76) |
| | Predicted no targeting peptide (noTP) | TargetP2.0 | |
| | Predicted mitochondrial targeting peptide (mTP) | TargetP2.0 | |

**Supplementary Table 7: Overall statistical summary of features across 12 included bacteria.**

| Feature & Subfeature | KS Statistic | KS p-value | AUROC |
|---|---|---|---|
| FEL mean_alpha | 0.078109 | 0.033649281 | 0.533735 |
| FEL mean_beta | 0.171354 | 6.49E-09 | 0.581166 |
| FEL mean_lrt | 0.123578 | 7.59E-05 | 0.448571 |
| FUBAR mean_alpha | 0.165362 | 2.39E-08 | 0.559793 |
| FUBAR mean_beta | 0.154702 | 2.31E-07 | 0.598928 |
| FUBAR mean_prob_neg | 0.069111 | 0.080625227 | 0.526963 |
| FUBAR mean_prob_pos | 0.043787 | 0.53372284 | 0.493673 |
| ProtLearn Amino acid composition | 0.005166 | 1 | 0.497654 |
| ProtLearn Average flexibility indices | 0 | 1 | 0.5 |
| ProtLearn Average volume of buried residue | 0.09418 | 9.30E-05 | 0.451399 |
| ProtLearn Composition | 0 | 1 | 0.5 |
| ProtLearn Hydropathy index | 0.007366 | 1 | 0.495871 |
| ProtLearn Hydrophobicity | 0.009006 | 1 | 0.495497 |
| ProtLearn Hydrophobicity index | 0.015681 | 0.998774961 | 0.507841 |
| ProtLearn Polarity | 0.015371 | 0.999113271 | 0.499691 |
| ProtLearn Relative mutability | 0.085558 | 0.000529226 | 0.551462 |
| ProtLearn Volume | 0.094731 | 8.28E-05 | 0.446842 |
| ProtLearn length | 0.184553 | 4.55E-17 | 0.61798 |
| ProtLearn pK (-COOH) | 0.00447 | 1 | 0.502235 |
| SLAC mean_dN/dS | 0.058973 | 0.1943912 | 0.492387 |
| SLAC mean_n | 0.081181 | 0.024674274 | 0.490426 |
| SLAC mean_s | 0.066445 | 0.102088656 | 0.477368 |
| allergenicity blast_score | 0.23513 | 6.10E-206 | 0.617828 |
| allergenicity hybrid_score | 0.241662 | 1.13E-217 | 0.642406 |
| allergenicity merci_score | 0.061766 | 1.87E-14 | 0.530883 |
| bcell avg_peptide_length | 0.211532 | 6.52E-120 | 0.632299 |

| | | | |
|---|---|---|---|
| bcell num_peptides | 0.246675 | 2.34E-163 | 0.655922 |
| cluster num_clusters | 0.046062 | 0.064054207 | 0.523092 |
| cluster percent_identity/num_strain | 0.320359 | 1.03E-77 | 0.610467 |
| deeplocpro prob_cell_wall_surface | 0.260574 | 2.07E-251 | 0.657703 |
| deeplocpro prob_cytoplasmic | 0.435443 | 0 | 0.242493 |
| deeplocpro prob_cytoplasmic_membrane | 0.266953 | 4.69E-264 | 0.643583 |
| deeplocpro prob_extracellular | 0.256569 | 1.23E-243 | 0.652091 |
| deeplocpro prob_outer_membrane | 0.244684 | 2.55E-221 | 0.660845 |
| deeplocpro prob_periplasmic | 0.183666 | 3.79E-124 | 0.596516 |
| deeptmhmm proportion_outside | 0.426839 | 0 | 0.725752 |
| discotope max_calibrated_score | 0.174844 | 7.39E-15 | 0.612061 |
| discotope max_discotope_score | 0.106609 | 8.67E-06 | 0.440914 |
| discotope max_rsa | 0.108449 | 5.64E-06 | 0.55841 |
| discotope mean_calibrated_score | 0.160946 | 1.19E-12 | 0.598475 |
| discotope mean_discotope_score | 0.149792 | 5.19E-11 | 0.406441 |
| discotope mean_rsa | 0.068005 | 0.012949023 | 0.543289 |
| discotope median_calibrated_score | 0.11178 | 2.55E-06 | 0.572559 |
| discotope median_discotope_score | 0.157067 | 4.57E-12 | 0.404465 |
| discotope median_rsa | 0.080478 | 0.001738262 | 0.543725 |
| discotope min_calibrated_score | 0.123591 | 1.25E-07 | 0.429925 |
| discotope min_discotope_score | 0.159335 | 2.09E-12 | 0.397854 |
| discotope min_rsa | 0.087222 | 0.000510093 | 0.453049 |
| discotope num_epitope_peptides | 0.202941 | 6.81E-20 | 0.614529 |
| dssp percent_helix | 0.10665 | 5.60E-06 | 0.480498 |
| dssp percent_loop | 0.06991 | 0.008106246 | 0.527561 |
| dssp percent_sheet | 0.1204 | 1.68E-07 | 0.50927 |
| ellipro mean_discontinuous_score | 0.106932 | 5.24E-06 | 0.563817 |
| ellipro mean_linear_score | 0.106932 | 5.24E-06 | 0.563817 |

| | | | |
|---|---|---|---|
| ellipro med_discontinuous_score | 0.065736 | 0.015302769 | 0.536742 |
| ellipro med_linear_score | 0.065736 | 0.015302769 | 0.536742 |
| ellipro num_discontinuous_epitopes | 0.133415 | 4.11E-09 | 0.5862 |
| ellipro num_linear_epitopes | 0.133415 | 4.11E-09 | 0.5862 |
| ifnepitope2 mean_blast_score | 0 | 1 | 0.5 |
| ifnepitope2 mean_ml_score | 0.038022 | 9.62E-06 | 0.481431 |
| ifnepitope2 mean_total_score | 0.038022 | 9.62E-06 | 0.481431 |
| ifnepitope2 med_blast_score | 0 | 1 | 0.5 |
| ifnepitope2 med_ml_score | 0.037014 | 1.82E-05 | 0.484065 |
| ifnepitope2 med_total_score | 0.037014 | 1.82E-05 | 0.484065 |
| mhci num_peptides | 0.179952 | 3.76E-120 | 0.603922 |
| mhci num_strong_binders | 0.180192 | 1.79E-120 | 0.601064 |
| mhci num_weak_binders | 0.177665 | 4.17E-117 | 0.6043 |
| mhci percentile | 0.050422 | 8.99E-10 | 0.518553 |
| mhci score | 0.042977 | 3.23E-07 | 0.488358 |
| mhcii num_peptides | 0.165666 | 2.00E-101 | 0.596859 |
| mhcii num_strong_binders | 0.151004 | 2.96E-84 | 0.588179 |
| mhcii num_weak_binders | 0.166352 | 2.86E-102 | 0.59717 |
| mhcii percentile | 0.04926 | 2.58E-09 | 0.490534 |
| mhcii score | 0.054396 | 2.89E-11 | 0.48399 |
| mixmhc2pred rank_best | 0.019626 | 0 | 0.512646 |
| rate4site max_per_site_score | 0.112121 | 3.42E-09 | 0.543433 |
| rate4site mean_per_site_score | 0.040116 | 0.148281209 | 0.504633 |
| rate4site median_per_site_score | 0.060554 | 0.005463399 | 0.52474 |
| rate4site min_per_site_score | 0.045204 | 0.073875333 | 0.503721 |
| rate4site_deeptmhmm max_outside_scores | 0.065129 | 8.86E-13 | 0.522742 |
| rate4site_deeptmhmm mean_outside_scores | 0.082978 | 1.77E-20 | 0.511845 |
| rate4site_deeptmhmm | 0.040436 | 3.43E-05 | 0.495918 |

| | | | |
|---|---|---|---|
| min_outside_scores | | | |
| signalp prob_other | 0.267012 | 1.93E-266 | 0.3312 |
| signalp prob_signalp | 0.267012 | 1.93E-266 | 0.6688 |
| targetp prob_SP | 0.294697 | 0 | 0.646759 |
| targetp prob_mTP | 0.042167 | 5.78E-07 | 0.512215 |
| targetp prob_noTP | 0.271514 | 1.21E-275 | 0.365693 |

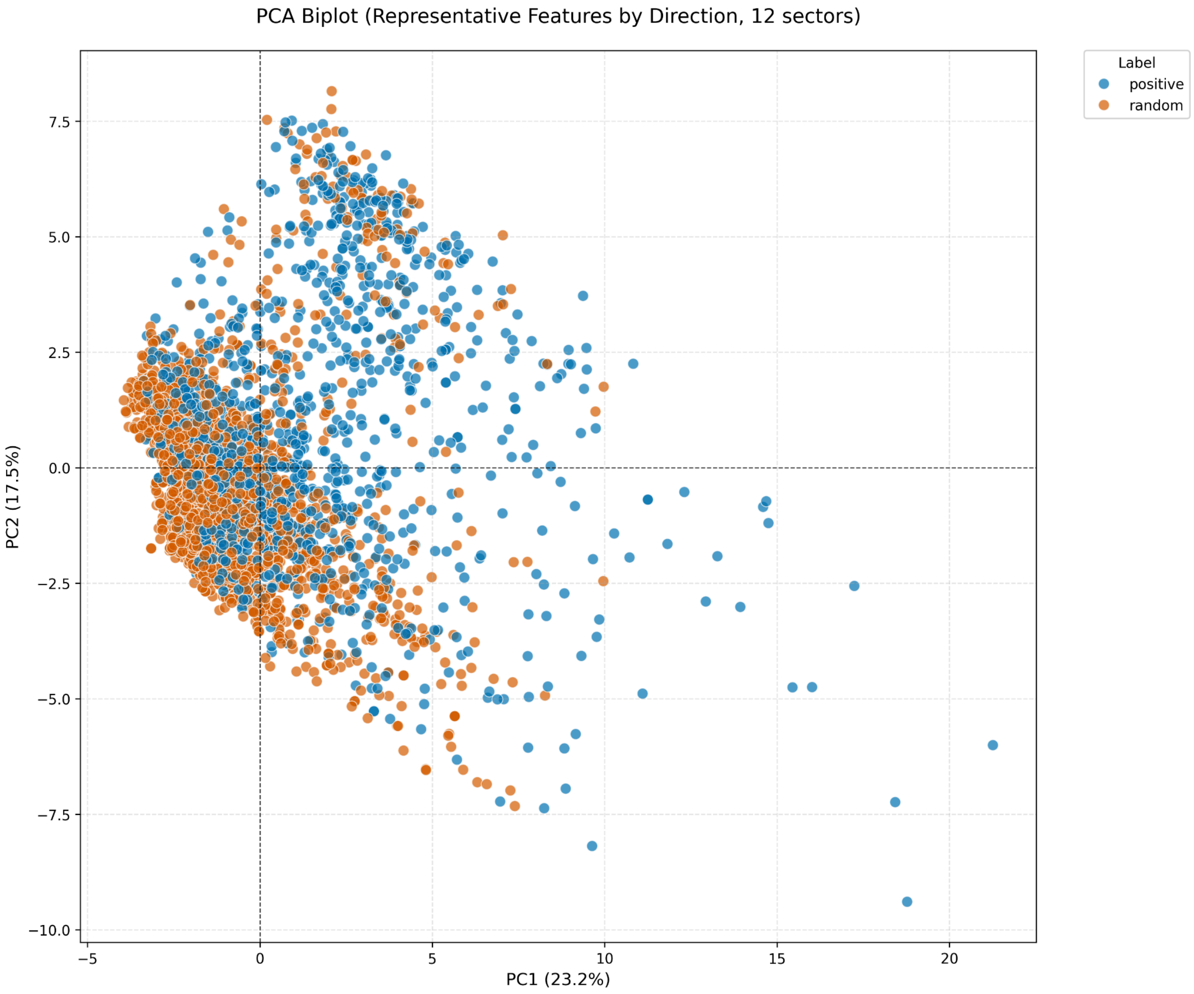


**Supplementary Figure 3: PCA biplot of all bacterial proteins based on PC1 and PC2.** PC1 and PC2 account for 23.2% and 17.5% of the total variance, respectively, explaining a combined 40.7% of the data variance. Each point represents a protein accession, categorized as antigenic proteins (blue) or non-antigenic proteins (vermillion).

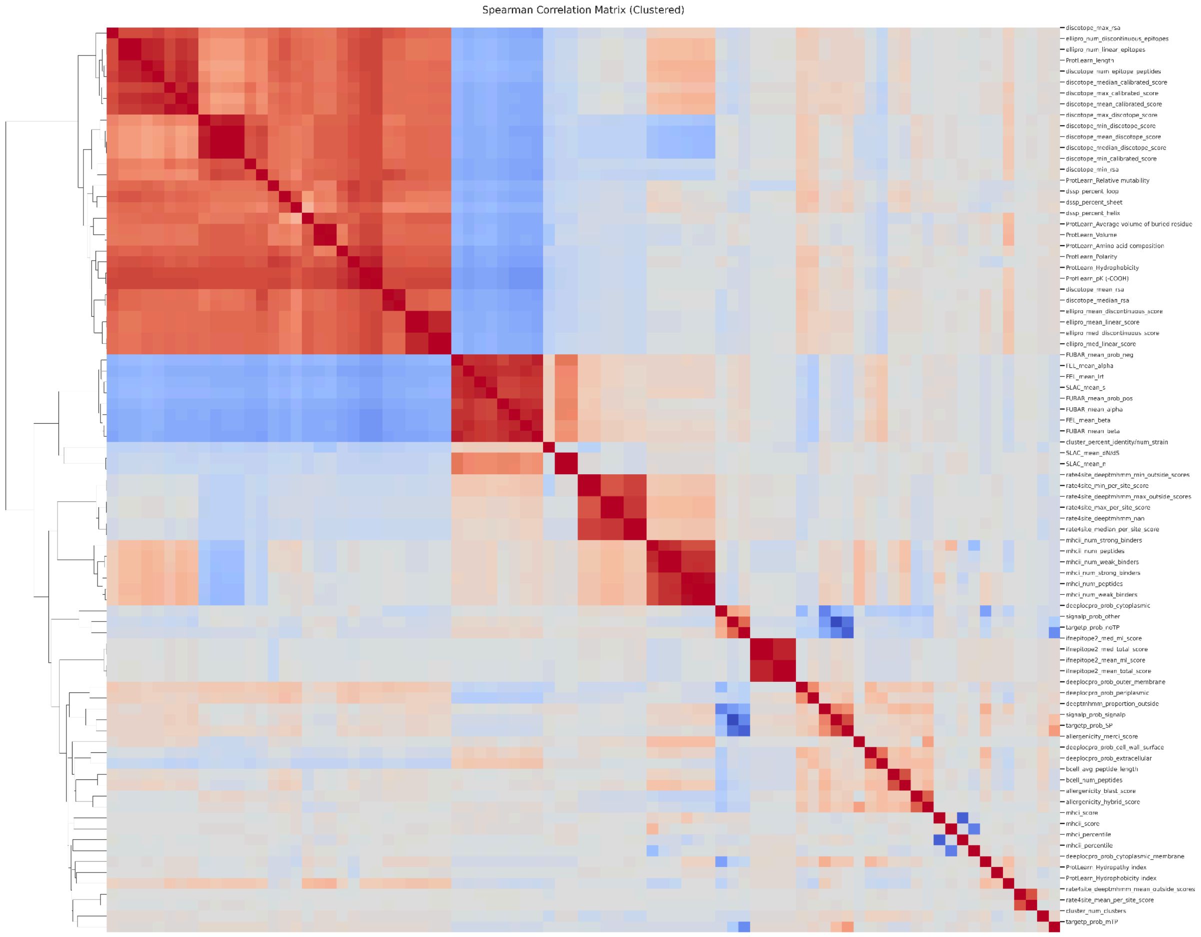


**Supplementary Figure 4: Spearman correlation matrix of all extracted protein features (clustered).** A clustered heatmap of the Spearman rank correlation coefficients between all extracted protein features. The color scale represents the strength and direction of correlations, with red indicating positive correlations and blue indicating negative correlations. Hierarchical clustering was applied to both rows and columns to group features with similar correlation profiles.

**Supplementary Table 8: Complete list of antigen predicted *Staphylococcus aureus* proteins and their antigen probability.** 210 out of 1002 proteins were filtered out due to allergenicity, 24 were filtered out due to high homology with human proteins, 133 were predicted to be non-antigens and were filtered out, and 374 proteins were collapsed as homologous duplicates. This list contains 308 *Staphylococcus aureus* predicted antigens. Four proteins (highlighted in red) were excluded as suitable *S. aureus* vaccine targets due to their prevalence across other bacterial species

| Accession(s) | Antigen Probability | Protein Name(s) |
|---|---|---|
| A0A3A5LJL8, Q6GHV7, A0A8B0LTH2, A0A0D6HSN6 | 0.734 | Fur-regulated protein B, Iron-regulated surface determinant protein B IsdB, Staphylococcal iron-regulated protein H, Staphylococcus aureus surface protein J |
| A0A390V9S3, A0AAJ5CTA4, A0A0D6GBZ3 | 0.728 | Deferrochelatase, Peroxidase EfeB |
| Q33C64 | 0.724 | Similar to staphylocoagulase |
| A0AAJ4YZF6, A0A389PZW7, A0A2C9TXC9, A0AAX2MNS4, A0A641A9A1, A0AAJ4YXI6, A0A2Y1RIG2, A0A0E8JWI2 | 0.722 | Staphopain A, Cysteine protease SspB, Cysteine protease, Staphopain B, Glycoside hydrolase family protein |
| A0A166JT38 | 0.718 | Exotoxin |
| X5E203, A0A390B6L2, A0AAE2ZUT4, A0A166JMZ6 | 0.714 | ABC transporter, periplasmic spermidine putrescine-binding protein potD, Extracellular solute-binding protein, Spermidine, putrescine ABC transporter substrate-binding protein, putrescine ABC superfamily ATP binding cassette transporter, binding protein |
| O54461 | 0.712 | Ampicillin resistance protein, Beta-lactamase, DUF4888 domain-containing protein, Phage related secreted protein Ear (Superantigen-encoding pathogenicity islands SaPI) |
| A0AAJ4YZ19, W8UY38 | 0.71 | Micrococcal nuclease, Staphylococcal nuclease, Thermonuclease |
| A0AAJ5CS40 | 0.71 | Oligopeptide transporter putative substrate binding domain protein |
| A0A389UFR6, A0A641A8N9, Q2FVE7, A0AAE2ZWA6 | 0.708 | Oligopeptide ABC superfamily ATP binding cassette transporter, binding protein, Staphylopine-dependent metal ABC transporter substrate-binding protein CntA, Metal-staphylopine-binding protein CntA, Oligopeptide transporter putative substrate binding domain protein |
| A0A113W024 | 0.708 | Exotoxin, Superantigen-like protein |
| A0AAE8PBX8, Q9S674, | 0.708 | Protein DltD |

| A0A9P3DMJ7 | | |
| --- | --- | --- |
| Q2G0X7, A0AAE2ZX53 | 0.704 | Staphylococcal superantigen-like 3, Exotoxin, Superantigen-like protein SSL3 |
| A0AAE8TIF0, A0A166IYH7, A0A391DGH4, A0A0D3Q5Z8, A0A641AA31, A0A389VEG0, A0AAE2ZV74 | 0.702 | Oligopeptide ABC transporter substrate-binding protein, Oligopeptide-binding protein oppA, Oligopeptide ABC transporter, periplasmic oligopeptide-binding protein oppA, Peptide ABC transporter substrate-binding protein, Oligopeptide ABC superfamily ATP binding cassette transporter, binding protein |
| A0A0D6GBP6 | 0.702 | Superantigen-like protein |
| A0A389VE54 | 0.7 | Superantigen-like protein |
| A0A166Q7U6 | 0.7 | Fibrinogen-binding protein |
| A0A641A9N7 | 0.698 | Fibrinogen-binding protein |
| A0AAJ5CSI3 | 0.698 | Micrococcal nuclease, Staphylococcal nuclease, Thermonuclease |
| P0A074, A0AAE2ZUJ5, Q6GE14 | 0.696 | Gamma-hemolysin component A, H-gamma-2, H-gamma-II, Bi-component gamma-hemolysin HlgAB subunit A |
| A0A166LCE3, A0A5C8X6J1, A0A389U9A3 | 0.696 | ABC-type nitrate, sulfonate, bicarbonate transporter, TauA, ABC transporter substrate-binding protein |
| A0A391ERX3, A0AAJ4YXY2, A0A0E8J2S9, A0AAE8PB46 | 0.696 | SceD-like transglycosylase, biomarker for vancomycin-intermediate strains, Transglycosylase, Transglycosylase SceD, Lytic transglycosylase SceD |
| A0A641A9C1 | 0.696 | 5'-nucleotidase, lipoprotein e(P4) family |
| A0A162H328 | 0.694 | Immunoglobulin-binding protein Sbi |
| A0A9P3DMY9 | 0.692 | Fibrinogen-binding protein |
| A0AAX2JXW1, A0AAJ4YW13, Q6GB45, Q2G015 | 0.692 | Clumping factor, Clumping factor A, Fibrinogen receptor A, Fibrinogen-binding protein A |
| A0AAJ4YZ87 | 0.692 | Exotoxin |
| A0A389XS11 | 0.69 | Immunoglobulin-binding protein Sbi |
| A0A224B3G8 | 0.69 | Staphylokinase |
| A0A166J4L2 | 0.69 | Exotoxin |
| A0A9P3DML6 | 0.688 | Staphylokinase |

| | | |
|---|---|---|
| A0AAX2MKD2 | 0.688 | Exotoxin |
| A0A3A5M257 | 0.688 | Exotoxin 15, Superantigen-like protein |
| A0AAE8PBT4, A0A0D1JUU6 | 0.686 | Autolysin, adhesin Aaa, N-acetylmuramoyl-L-alanine amidase |
| A0A6B2IZR7 | 0.686 | Oligopeptide ABC transporter substrate-binding protein |
| P02976 | 0.686 | Immunoglobulin G-binding protein A, Staphylococcal protein A |
| A0AAX2MI48 | 0.686 | Secreted von Willebrand factor-binding protein, Staphylocoagulase |
| Q2FZR2 | 0.686 | Oligopeptide ABC transporter, substrate-binding protein, putative |
| P0C1U7, A0AAE2ZTH7, W8U332, A0A1J0J3G8, A0AAJ4YV78, A0A0D6WBZ7, A0A2S6D3A1, A0AAE2ZU90, A0A518QF20, A0A166LCU2, W8TR90, W8USR6, X5E526, A0AAE3D3N5, A0A2S6DCI1, A0AAJ4YW98, A0A169E8V2, A0A2U0IJR9, A0A641A6Q5, A0AAJ4YX04, W8U7V9, A0A3A2IVX8, A0A641A7K4, Q6GJK9 | 0.684 | N-acetylmuramoyl-L-alanine amidase sle1, CHAP domain-containing protein, N-acetylmuramoyl-L-alanine amidase, Peptidase M23B, Secretory antigen SsaA, Staphyloxanthin biosynthesis protein, Secretory antigen, LysM domain protein, LysM peptidoglycan-binding domain-containing protein, Peptidase M23, Peptidoglycan hydrolysis related protein |
| A0A0D3Q4R4, A0A090N182, A0A641AA78, A0A7I0YLT7, A0A2S6D767, A0A390BCG4, A0AAE2ZW72, B0I1W0, A0A166DEB6, A0A3A5LS85 | 0.684 | Exported protein, Uncharacterized protein conserved in bacteria, YxeA family protein, DUF1093-containing protein, Putative DUF1093-containing protein |
| C3VIQ5 | 0.684 | Staphylokinase |
| P0A015 | 0.684 | Immunoglobulin G-binding protein A, Staphylococcal protein A |

| | | |
|---|---|---|
| A0AAE8PBB6 | 0.682 | Exotoxin, Superantigen-like protein SSL14 |
| A0A0D6HSA5 | 0.682 | Exotoxin, Superantigen-like protein |
| A0AAX2MMZ9 | 0.682 | Staphylokinase |
| Q2G1S8, A0A641A8M2 | 0.682 | Staphylococcal superantigen-like 4, Superantigen-like protein SSL4 |
| P0C6P2 | 0.68 | Fibrinogen-binding protein |
| E2D9B8, F8WKG6 | 0.678 | PBP2a family beta-lactam-resistant peptidoglycan transpeptidase MecA, Penicillin-binding protein 2 |
| C4B833 | 0.678 | Staphylocoagulase |
| B2NJ60 | 0.676 | Coagulase |
| A0AAX2K0P1 | 0.676 | Uncharacterized protein |
| B6VQW5 | 0.674 | Staphylocoagulase |
| A0A162GR70 | 0.672 | Exotoxin |
| A0A2C9TVL9 | 0.672 | ABC transporter substrate-binding protein |
| E2GAH4 | 0.672 | von Willebrand factor-binding protein |
| A0A659IBA1 | 0.672 | Exotoxin, Superantigen-like protein SSL14 |
| A0A0D6GBH7 | 0.67 | Superantigen-like protein |
| A0AAJ4YZC1 | 0.67 | Lipoprotein |
| C4B818 | 0.67 | Staphylocoagulase |
| A0AAJ4YX07, Q9RQG6, A0AAX2JYZ4 | 0.668 | RND multidrug efflux transporter Acriflavin resistance protein, Efflux RND transporter permease subunit, MMPL family transporter |
| A0AAE2ZU14, A0A641AAM4 | 0.666 | Exotoxin, Superantigen-like protein SSL2 |
| A0AAX2JYJ2, A0AAE3D4M0, A0A9P2Z0D5, Q2G0L4, Q6GBS5, A0A9P3DMH0, Q8NXX5, A0A9P2YY90, A0AAJ4YWK2 | 0.666 | SdrE protein, SdrH family protein, SdrH protein, Ser-Asp rich fibrinogen, bone sialoprotein-binding protein SdrH, Serine-aspartate repeat-containing protein D, MSCRAMM family adhesin SdrD, bone sialoprotein-binding protein SdrD, Serine-aspartate repeat-containing protein E, bone sialoprotein-binding protein SdrE, Ser-Asp rich fibrinogen-binding, bone sialoprotein-binding protein |
| A0AAJ4YVZ0 | 0.666 | Lipoprotein |
| A0A0D3Q6S2 | 0.666 | Fibrinogen-binding protein |

| | | |
|---|---|---|
| A0AAE8TJK5 | 0.666 | Alpha-hemolysin, Alpha-toxin |
| A0A641A7L2 | 0.664 | Micrococcal nuclease, Staphylococcal nuclease, Thermonuclease |
| A0AAJ4YZA4 | 0.664 | Exotoxin |
| A0A641A8Q3 | 0.664 | Superantigen-like protein SSL11 |
| A0A162G2W5 | 0.664 | Lipoprotein |
| A0A0D3Q462 | 0.664 | Uncharacterized protein |
| A0AAE8TJ59 | 0.662 | Micrococcal nuclease, Staphylococcal nuclease, Thermonuclease |
| A0A389V673 | 0.662 | Uncharacterized protein |
| A0A641AAA6 | 0.66 | Uncharacterized protein |
| Q9LC46 | 0.66 | Staphylokinase |
| X5E2Q9 | 0.66 | Lipoprotein, Putative lipoprotein |
| A0A380DP26 | 0.66 | Exotoxin, Superantigen-like protein |
| A0A3A2ZYW2 | 0.66 | Micrococcal nuclease, Staphylococcal nuclease, Thermonuclease |
| A0A0D3QAF5 | 0.658 | Immunoglobulin-binding protein Sbi |
| A0AAE8PBB7, W8UTZ7, A0A166NTM9 | 0.658 | Maltodextrin-binding protein |
| A0A641AB29 | 0.658 | Superantigen-like protein SSL6 |
| A0A3A5LNY2 | 0.658 | Uncharacterized protein |
| A0AAJ5CU77 | 0.658 | Leukocidin F subunit |
| X5DYK9 | 0.656 | Exotoxin, Superantigen-like protein |
| A0A0D6H7X3, A0AAJ4YVQ1, A0A389XYY8 | 0.656 | Hyaluronate lyase, Polysaccharide lyase 8 family protein |
| A0A641A3T5 | 0.656 | Cysteine protease staphopain A |
| A0AAJ4YV91 | 0.656 | Lipoprotein |
| A0A0D6GBK0, A0A3M9H3Z7 | 0.654 | Exotoxin, Superantigen-like protein |
| A0A0Z0MT78 | 0.654 | Hyaluronate lyase |
| A0AAE8PDG0, A0A389UE07 | 0.654 | Exotoxin, Superantigen-like protein SSL13, Superantigen-like protein |

| | | |
|---|---|---|
| Q9S446, A0A4P7PAF7, A0AAE8PBZ9, A0A0M0Z384 | 0.652 | Sortase, Sortase A, Class A sortase SrtA, LPXTG specific sortase A, Sortase A transpeptidase |
| A0A0D6GCF3 | 0.652 | Enterotoxin-like toxin X |
| A0A2Y1TBB7, Q2FC27, Q1XG18, P00807 | 0.652 | Beta-lactamase, Penicillinase |
| Q2FWN1 | 0.652 | Membrane anchored protein |
| A0A0D3Q4F9 | 0.652 | Exotoxin, Superantigen-like protein |
| A0A0D6I075 | 0.652 | Micrococcal nuclease, Staphylococcal nuclease, Thermonuclease |
| B1Q017 | 0.65 | Gamma-hemolysin subunit A, LukS-PV, Panton-Valentine bi-component leukocidin subunit S, Panton-Valentine leucocidin S, Panton-valentine Leukocidin S |
| A0AAE8P9H1 | 0.65 | Hyaluronate lyase, Polysaccharide lyase 8 family protein |
| A0A391JVV4 | 0.65 | Efflux RND transporter periplasmic adaptor subunit, Membrane-fusion protein-like protein |
| A0A166E754 | 0.65 | Membrane protein, Protein of uncharacterized function (DUF1433) |
| A0AAE2ZU05 | 0.65 | Tandem-type lipoprotein |
| X5DSM8 | 0.65 | Superantigen-like protein |
| A0A1D4H1Q1 | 0.65 | DUF5067 domain-containing protein, Putative lipoprotein |
| A0A390AFP1 | 0.65 | Superantigen-like protein |
| A0AAE8P9U8, A0A166PFD1, Q70MK4, A0A0D6H3T6 | 0.65 | Molybdate ABC transporter substrate-binding protein, Molybdate-binding protein, Molybdenum ABC transporter substrate-binding protein, Molybdenum (Mo2+) ABC superfamily ATP binding cassette transporter, binding protein |
| A0AAJ4YYQ0, W8U629 | 0.648 | Phosphate-binding protein |
| A0A7U7ET86 | 0.648 | Immunoglobulin G-binding protein A, Staphylococcal protein A |
| W8UXH5 | 0.648 | Lipoprotein |
| A0AAJ5CTI4 | 0.648 | Membrane lipoprotein |
| A0A131JTH8 | 0.648 | Uncharacterized protein |
| Q6G6Q1, A0A166QIC8, A0A3M7ZM78, | 0.648 | Gamma-hemolysin component C, Gamma-hemolysin subunit A, Leukocidin S subunit, Bi-component gamma-hemolysin HlgCB subunit C, HlgC, H-gamma-1, H-gamma-I |

| | | |
|---|---|---|
| Q07227, A0AAE8P9W0 | | |
| A0A390ZI89 | 0.648 | DUF1433 domain-containing protein, Protein of uncharacterized function (DUF1433) |
| A0A641A8U6 | 0.648 | Immunoglobulin-binding protein Sbi |
| A0AAE8PD83 | 0.648 | Enterotoxin-like toxin X |
| A0A1S6D4V8, A0A218M1Q9, A0AAJ5CSS1, A0A0Z4LY14 | 0.646 | D-alanyl-D-alanine carboxypeptidase, DUF1958 domain-containing protein, Penicillin binding protein 4, Penicillin-binding protein PBP4, Serine-type D-Ala-D-Ala carboxypeptidase, Penicillin binding protein PBP4 |
| A0AAJ5CSE5 | 0.646 | Lipoprotein |
| A0AAE8TK97, A0A390DBJ6, A0A0D6HCK9 | 0.646 | Bi-component leukocidin LukGH subunit G, Gamma-hemolysin component B, Leukocidin, hemolysin toxin family protein, Gamma-hemolysin subunit B |
| A0A3A5LL43 | 0.646 | AcrB/AcrD/AcrF family protein, Efflux RND transporter permease subunit, RND superfamily resistance-nodulation-cell division acriflavin:proton (H+) antiporter |
| A0AAQ1CIB4 | 0.644 | N-acetylmuramoyl-L-alanine amidase |
| A0A2I7YBL3, A0A641A666 | 0.644 | Sensor protein kinase WalK |
| A0A2U0IK27 | 0.644 | Membrane lipoprotein, Tandem-type lipoprotein |
| A0A0D1IUM4 | 0.644 | Exported protein |
| A0A3A2ISC3 | 0.644 | Exotoxin, Superantigen-like protein |
| A0A2C9TYK2 | 0.644 | Putative lipoprotein, Staphylococcal tandem lipoprotein, Tandem-type lipoprotein Lpl9 |
| A0A166MVX8 | 0.644 | Lipoprotein, Uncharacterized protein conserved in bacteria |
| W8TTS4 | 0.642 | C39 family peptidase, Lipoprotein, Uncharacterized protein conserved in bacteria |
| A0A5N5IQ84 | 0.642 | Tandem-type lipoprotein |
| A0A391DPH1 | 0.642 | Exotoxin, Superantigen-like protein |
| A0A0D6GXZ6 | 0.642 | DUF1433 domain-containing protein, Protein of uncharacterized function (DUF1433) |
| C4B852 | 0.64 | Staphylocoagulase |
| A0A641A7H5 | 0.64 | Amidase domain-containing protein, N-acetylmuramoyl-L-alanine amidase, family 4 |

| | | |
|---|---|---|
| A0AAJ4YWH1, A0AAE2ZVK6 | 0.64 | ABC superfamily ATP binding cassette transporter, binding protein, ABC transporter substrate-binding protein |
| Q6GJM8 | 0.64 | Uncharacterized lipoprotein SAR0445 |
| A0AAJ4Z045, A0A1M4RHY1, A0A5D3UEU8, A0A2X3QG95, A0AAE8TJU3 | 0.64 | Biofilm polysaccharide intercellular adhesin deacetylase, Intercellular adhesion protein B, Poly-beta-1, 6-N-acetyl-D-glucosamine N-deacetylase |
| A0A0U1MMC4, W8UA89, A0AAJ4YUY7 | 0.638 | Multifunctional fusion protein |
| A0A389U7S2 | 0.638 | Amidase domain-containing protein, N-acetylmuramoyl-L-alanine amidase, family 4 |
| A0AAE2ZVL7 | 0.636 | Immunoglobulin-binding protein Sbi |
| A0A6K1K184 | 0.636 | ABC superfamily ATP binding cassette transporter, binding protein, ABC transporter substrate-binding protein |
| A0A391A4D9 | 0.636 | CHIPS, Chemotaxis inhibitory protein |
| A0AAJ5CTZ3 | 0.636 | Membrane anchored protein |
| A0AAX2K1M1 | 0.634 | Lipoprotein |
| A0AAJ5CTM9 | 0.634 | Staphylococcal exotoxin |
| A0A641A684 | 0.634 | Bi-component leukocidin LukGH subunit G, Gamma-hemolysin component B |
| A0A162FEP7, A0A3A3ACW7, X5EIX3, A0AAE8PAR1, A0A390B8Q0 | 0.634 | PTS glucose EIICBA component, PTS glucose transporter subunit IICBA, PTS system transporter subunit IIABC, PTS transporter subunit EIIC, PTS system, glucose-specific IICBA component, PTS glucose transporter subunit IIB, PTS system glucose-specific transporter subunit IIBC, PTS system maltose, glucose-specific IICB component |
| A0A2W3WBR7 | 0.632 | Lipoprotein |
| A0A181GKX6 | 0.632 | Penicillin binding protein 4 |
| A0A641AC27 | 0.632 | Superantigen-like protein SSL12 |
| A0A641A990 | 0.632 | Enterotoxin-like toxin X |
| Q57227, A0AAE8TIU7, P0A077 | 0.632 | F component of leucocodin R, Gamma-hemolysin component B, Gamma-hemolysin subunit B, HlgB-like ORF protein, Bi-component gamma-hemolysin HlgAB, HlgCB subunit B, H-gamma-1, H-gamma-I |
| A0A391F9N7, A0A641A9N0 | 0.632 | Staphylococcal tandem lipoprotein, Tandem lipoprotein within Pathogenicity island, Tandem-type lipoprotein, Tandem-type |

| | | |
|---|---|---|
| | | lipoprotein Lpl5 |
| A0A641A7R9 | 0.632 | Hyaluronate lyase |
| A0A391EL89 | 0.63 | Enterotoxin-like toxin X |
| A0A2W3TG75 | 0.628 | Fibrinogen-binding protein |
| A0AAJ5CTK1 | 0.628 | Staphylococcal tandem lipoprotein |
| A0A641A7Q8 | 0.628 | Tandem-type lipoprotein |
| A0A391EKT5 | 0.628 | Staphylococcal tandem lipoprotein, Tandem-type lipoprotein |
| A0A0Y9RFD4 | 0.628 | Amidase domain-containing protein, N-acetylmuramoyl-L-alanine amidase, family 4 |
| A0AAE8PDZ2 | 0.628 | Amidase domain-containing protein, N-acetylmuramoyl-L-alanine amidase, family 4 |
| A0AAE8PDQ6 | 0.628 | Exotoxin, Superantigen-like protein SSL12 |
| A0A2U0IPK5 | 0.626 | Tandem-type lipoprotein |
| X5DZ18, A0A3A1VX69, A0A131JPX7 | 0.626 | ESAT-6 secretion machinery protein EssA |
| A0A389QCM1 | 0.626 | Staphylococcal tandem lipoprotein, Tandem-type lipoprotein |
| A0A641ABA7 | 0.626 | ABC transporter substrate-binding protein |
| A0A641A8J4 | 0.624 | NDxxF motif lipoprotein |
| A0AAJ5CTN4 | 0.624 | Staphylococcal tandem lipoprotein |
| A0A641A9D2 | 0.624 | Tandem-type lipoprotein |
| A0A068A6G0 | 0.624 | CHIPS, Chemotaxis inhibitory protein |
| A0A0D6WII1 | 0.624 | Alpha-hemolysin, Alpha-toxin |
| A0A0U1MK78 | 0.624 | Fibrinogen-binding protein |
| A0AAJ4YUB2, X5DW82, A0A4U2KP01 | 0.622 | SPP1 family prophage L54a, Holin, Phage holin |
| A0AAJ5CSE2 | 0.622 | Extracellular matrix and plasma binding protein |
| A0A2L2E298 | 0.622 | Putative lipoprotein |
| A0AAE3D6P3 | 0.622 | DUF1433 domain-containing protein, Membrane protein |
| A0A641A6Y4 | 0.622 | Tandem-type lipoprotein |
| A0A2C6MVP5 | 0.622 | Beta-channel forming cytolysin, Gamma-hemolysin subunit A, |

| | | |
|---|---|---|
| | | Leukocidin S subunit lukE |
| A0A390B4P9 | 0.622 | Lipoprotein, Tandem-type lipoprotein |
| A0A6B5F8B5 | 0.62 | Secreted protein |
| A0A391AB27 | 0.618 | Lipoprotein |
| A0A641A7L3 | 0.618 | Efflux RND transporter permease subunit |
| A0A0D6GCD0 | 0.616 | Lipoprotein, PepSY domain-containing protein, Peptidase |
| A0A389VCE2 | 0.616 | DUF5067 domain-containing protein, Lipoprotein |
| Q2G0L5, Q6GBS6, A0AAX2JYB1, Q6GJA7, Q7A781 | 0.614 | Serine-aspartate repeat-containing protein C, Surface anchored protein |
| A0AAE8TKH1 | 0.612 | Lipoprotein |
| A0A0D3Q6S7 | 0.612 | Alpha-hemolysin, Alpha-toxin |
| Q6SV31 | 0.612 | Alpha-hemolysin, Alpha-toxin |
| A0A266CTA0 | 0.61 | DUF1433 domain-containing protein, Protein of uncharacterized function (DUF1433) |
| A0A641A7W2 | 0.61 | Tandem-type lipoprotein |
| A0A1M3SHT7 | 0.61 | PTS glucose EIICBA component, PTS system glucose-specific transporter subunit IICBA, PTS system transporter subunit IIABC |
| A0AAX2MJ33 | 0.608 | PTS system transporter subunit IIABC |
| A0A4P7P3I1 | 0.608 | Tandem-type lipoprotein |
| A0AAJ5CS58 | 0.608 | Lipoprotein |
| A0AAE8PAW7 | 0.608 | DUF4467 domain-containing protein, Lipoprotein |
| A0A0D6HDR3 | 0.608 | Lipoprotein |
| A0A389PZU7, A0A0D6HC73, A0AAE8PCH5 | 0.608 | Bi-component leukocidin LukGH subunit H, Leukocidin S subunit, Leukocidin, lemolysin toxin family protein, Succinyl-diaminopimelate desuccinylase |
| Q6GJN3 | 0.606 | Uncharacterized lipoprotein SAR0438 |
| A0A641A4M0 | 0.606 | Uncharacterized protein |
| A0A0D3QAH7 | 0.604 | DUF1433 domain-containing protein, Membrane protein, Protein of uncharacterized function (DUF1433) |
| A0A133Q1H1 | 0.604 | Calcium-binding protein, Excalibur calcium-binding domain-containing protein |

| | | |
|---|---|---|
| A0A2C6MZ76 | 0.604 | Tandem-type lipoprotein |
| A0A2C9TL89 | 0.604 | Cystatin-like fold lipoprotein, DUF4467 domain-containing protein |
| A0A641A5G3 | 0.604 | Calcium-binding protein |
| W8U6L2 | 0.602 | Lipoprotein, PepSY domain-containing protein, Peptidase, Secreted protease inhibitor |
| A0A0E1XCS8 | 0.602 | Tandem lipoprotein |
| A0A8A7XIF5 | 0.602 | CAP domain-containing protein |
| A0AAE3D5Q0 | 0.602 | CAP domain-containing protein |
| A0A391E3Q8 | 0.602 | DUF1433 domain-containing protein, Membrane protein |
| X5E258 | 0.6 | Competence protein ComGE |
| A0A2Y9TLP4 | 0.598 | Lipoprotein |
| A0A641ABA5 | 0.598 | Tandem-type lipoprotein |
| A0A641A4E5 | 0.598 | Bi-component leukocidin LukGH subunit H |
| A0A6B5S7D5 | 0.596 | CHIPS, Chemotaxis inhibitory protein |
| W8U4T8 | 0.596 | Membrane spanning protein, Metal-dependent hydrolase |
| A0A266CN08 | 0.596 | Tandem-type lipoprotein |
| A0A641A8Q8 | 0.596 | Superantigen-like protein SSL8 |
| A0AAJ4YV34, Q9EYL5, A0A0D1HWZ3 | 0.596 | Protein-disulfide isomerase, DsbA-like protein, Disulfide bond protein A, DsbA family protein, Thioredoxin domain-containing protein, Disulfide bond formation protein D |
| X5DWN8 | 0.594 | M50 family metallopeptidase, M50 family peptidase, Membrane protein, Membrane spanning protein |
| A0A166J0L6 | 0.594 | Membrane spanning protein, Membrane-bound metal-dependent hydrolase |
| A0A3A3ALI8 | 0.594 | Lipoprotein, Tandem-type lipoprotein |
| A0A641A9G9 | 0.594 | Tandem-type lipoprotein |
| A0A166MWQ0 | 0.594 | Membrane protein |
| A0A6B5E444 | 0.594 | Tandem-type lipoprotein |
| A0A7U7EWK0 | 0.594 | Transport protein |
| W8U9Q7, W8UWJ4 | 0.592 | Exported protein, Gas vesicle protein, YtxH domain-containing protein, Membrane associated protein, Putative staphylococcal protein |

| | | |
|---|---|---|
| Q93UU8, A0A390W8I0, A0A1S5YLW8 | 0.592 | Bi-component leukocidin LukED subunit D, Leucotoxin LukD, Leukotoxin LukD, LukNF, Leukocidin LukD, Panton-Valentine bi-component leukocidin subunit F |
| A0A6B5TU88 | 0.592 | Tandem-type lipoprotein |
| A0A641A618 | 0.59 | DUF1433 domain-containing protein |
| A0A090LUS6, A0AAX2MNC3, X5EK72, A0A391FC21 | 0.59 | DUF1307 domain-containing protein, Lipoprotein |
| A0AB74Q154 | 0.59 | Membrane spanning protein |
| A0AAE8PEE6 | 0.59 | DUF1433 domain-containing protein, Exported protein |
| A0A166Q7R0 | 0.59 | Fibrinogen-binding protein |
| A0A9N8IH31 | 0.588 | Membrane lipoprotein, Staphylococcal tandem lipoprotein |
| A0A390U9J4 | 0.588 | Lipoprotein |
| A0A166ND87 | 0.588 | Exported protein |
| A0A2U0IMZ5 | 0.588 | Secreted protein |
| A0A641A912, A0A131JP41, A0A3A3ATI1, A0A2C9TV59, A0AAE8PBA3 | 0.588 | Autolysin LytM, Glycyl-glycine endopeptidase LytM |
| A0A641AA18, A0AAJ4YYG6, A0A0D6HUN3, A0AAE3D4V2, A0A3A5LQS0 | 0.586 | NPQTN specific sortase B, SrtB family sortase, Class B sortase, Sortase B transpeptidase |
| A0A641A8T8 | 0.586 | RND transporter |
| A0A2I7YBH8 | 0.586 | DUF1433 domain-containing protein |
| A0A3M8VBL4 | 0.586 | Fibrinogen-binding protein |
| W8TNQ9, Q8VQS9, A0A090LVV4 | 0.586 | ABC superfamily ATP binding cassette transporter, binding protein, ABC transporter substrate-binding protein, Metal ABC transporter substrate-binding protein, MntC, SitC, Manganese ABC transporter substrate-binding lipoprotein |
| A0A5S9C563 | 0.586 | Protein of uncharacterized function (DUF1433) |
| A0AAE8TKI7 | 0.584 | Lipoprotein, Tandem-type lipoprotein |
| A0AAJ5CU80 | 0.584 | YbbR-like family protein |

| | | |
|---|---|---|
| A0A641A5V3 | 0.584 | Uncharacterized protein |
| A0A3A3AS45 | 0.584 | Membrane lipoprotein, Staphylococcal tandem lipoprotein, Tandem-type lipoprotein |
| A0A2L2F4N6 | 0.584 | Uncharacterized protein |
| A0A1S6DSG8 | 0.584 | Tandem-type lipoprotein |
| A0A2I7YBH6 | 0.584 | DUF4909 domain-containing protein |
| A0A0Z4NNI4 | 0.584 | Exported protein, Lipoprotein, putative |
| Q6G6I7 | 0.582 | Uncharacterized protein SAS2374 |
| Q9ZAI5 | 0.58 | CdaA regulatory protein CdaR, Uncharacterized protein orf2, YbbR-like domain-containing protein, YbbR-like family protein |
| A0A389Q5J4 | 0.58 | Lipoprotein, Tandem-type lipoprotein |
| A0AAJ5CU40 | 0.58 | Leukocidin S subunit |
| A0A2Y9TQ37 | 0.58 | Uncharacterized protein |
| A0AAJ5CSZ5 | 0.578 | Lipoprotein |
| A0A166N8V6 | 0.578 | Putative lipoprotein, Staphylococcal tandem lipoprotein |
| A0A2U0IFN2 | 0.578 | Calcium-binding protein |
| A0A9P2YX88 | 0.576 | Lipoprotein |
| A0AAN1ZDM6 | 0.576 | Transposon-related protein |
| A0A2U0IPJ7 | 0.576 | Tandem-type lipoprotein |
| A0A391DHU9 | 0.576 | Anaerobic ribonucleoside-triphosphate reductase activating protein, von Willebrand binding protein |
| A0A166M720 | 0.574 | Lipoprotein, ZK354.3 domain protein |
| A0AAX2MMQ3 | 0.574 | Staphylococcal tandem lipoprotein |
| X5EG17 | 0.574 | Lipoprotein, ZK354.3 domain protein |
| A0AAE2ZZ67 | 0.572 | Tandem-type lipoprotein |
| B6V385 | 0.572 | Putative exported protein, Transposon-related protein |
| A0AAE8PAQ7 | 0.572 | DUF4467 domain-containing protein, Lipoprotein |
| A0A0D3Q6F6 | 0.572 | Fibrinogen-binding protein |
| Q2G177 | 0.572 | DUF5079 family protein |
| A0A513Q6V6 | 0.568 | Sphingomyelin phosphodiesterase/beta-hemolysin |

| | | |
|---|---|---|
| A0A166JHB2 | 0.566 | Lipoprotein, putative, Protein of uncharacterized function (DUF1433) |
| A0AAE2ZYH5 | 0.566 | Tandem-type lipoprotein |
| A0AAE2ZSL6 | 0.566 | Tandem-type lipoprotein |
| A0AAE2ZRT7 | 0.566 | ZK354.3 domain protein |
| A0A2U0IFR6 | 0.566 | DUF4909 domain-containing protein |
| A0A0D1HKW8 | 0.564 | Lipoprotein, putative |
| A0A2W2JKA3 | 0.564 | M50 family metallopeptidase, M50 family peptidase, Membrane spanning protein |
| A0A0E1X437 | 0.564 | Excalibur domain protein |
| A0A5N5IGG7 | 0.562 | Tandem-type lipoprotein |
| A0A6M1ABI1 | 0.56 | Phage protein, Phi PVL-like protein |
| A0A7R6NZL1 | 0.56 | Late competence protein ComGE |
| A0A2U0IF80 | 0.56 | Tandem-type lipoprotein Lpl1 |
| A0AAE8PDC6 | 0.556 | Lipoprotein, Tandem-type lipoprotein |
| A0A641A794 | 0.554 | Uncharacterized protein |
| A0AAE8P9H8 | 0.554 | Lipoprotein |
| A0AAN1ZDM5 | 0.552 | Transposon-related protein |
| A0A391A4W5 | 0.55 | Lipoprotein |
| A0A0D6H001 | 0.55 | Lipoprotein |
| A0A641A5U9 | 0.548 | DUF4909 domain-containing protein |
| A0A389NBE8 | 0.546 | Lipoprotein |
| A0AAJ5CSZ9 | 0.546 | Transposon-related protein |
| Q2FVC5 | 0.546 | Uncharacterized lipoprotein SAOUHSC_02788 |
| A0A0D3QB23 | 0.542 | Lipoprotein |
| W8U485, A0AAJ4YXX9 | 0.54 | Foldase YidC, Membrane integrase YidC, Membrane protein YidC, Membrane protein insertase YidC |
| A0AAE8TJU8 | 0.54 | 1-phosphatidylinositol phosphodiesterase, Phosphatidylinositol diacylglycerol-lyase, Phosphatidylinositol-specific phospholipase C |
| A0AAE8TIR2 | 0.54 | Anaerobic ribonucleoside-triphosphate reductase activating protein, von Willebrand binding protein |

| | | |
|---|---|---|
| A0A641A9B3 | 0.54 | DUF5079 family protein |
| B6V384 | 0.534 | Lipoprotein, Lipoprotein, putative |
| A0AAE8TL59 | 0.534 | Lipoprotein |
| A0A114AKY3 | 0.534 | 1-phosphatidylinositol phosphodiesterase, Phosphatidylinositol diacylglycerol-lyase, Phosphatidylinositol-specific phospholipase C |
| A0A390WCR0 | 0.532 | DUF5079 family protein, Lipoprotein, putative |
| A0A115HMG3, W8TW02 | 0.53 | ComG operon protein 3 |
| Q7DIE8 | 0.528 | Delta-hemolysin, Delta-toxin |
| A0A0D6GL09 | 0.528 | 1-phosphatidylinositol phosphodiesterase, Phosphatidylinositol diacylglycerol-lyase, Phosphatidylinositol-specific phospholipase C |
| A0AAJ4YX42 | 0.528 | Lipoprotein, putative |
| A0A390W179 | 0.526 | Lipoprotein, Tandem-type lipoprotein |
| A0A9N8NS69 | 0.522 | 1-phosphatidylinositol phosphodiesterase, Phosphatidylinositol diacylglycerol-lyase, Phosphatidylinositol-specific phospholipase C |
| A0AAJ5CTW5 | 0.514 | Uncharacterized protein |
| W8TWS4 | 0.506 | Ribonuclease Y |

**Supplementary Table 9: Predicted *S. aureus* antigens that were removed due to allergenicity (AlgPred2 hybrid score > 0.3).** A total of 240 proteins collapsed into 205 rows.

| Accession(s) | Antigen Probability | Protein Names |
|---|---|---|
| P81177, A0A641ABP3, A0A391LQP8, A0A641A6Z5, A0A0D3QA93, A0A122DF78, A0A1K8I4B4, A0AAE2ZUL3, A0A0D1JPY7 | 0.651778 | Staphylococcus aureus neutral proteinase, Zinc metalloproteinase aureolysin, Zinc metallopeptidase, ABC superfamily ATP binding cassette transporter, binding protein, Zinc ABC transporter substrate-binding lipoprotein AdcA, Zn-binding lipoprotein adcA-like protein, Zinc ABC transporter substrate-binding protein |
| A0A2C9TSC4 | 0.76 | DNA-binding protein |
| A8QKE0 | 0.756 | Neutral metalloproteinase |
| A0A166DWI7 | 0.75 | ABC-type transporter, DNA-binding protein |
| A0AAE8TKW4, A0A9P2YYD8, A0A131JSY8, A0A8G2M8L9, A0AAX2K353, A0A389UCX7, Q9RQG6, A0AAQ1CLX2, X5EJN4, A0A6B5L736, Q7DI48, A0A9Q8DF91, A0A0D1H9A1, A0AAE2ZWD6, X5E0B5, A0A166N5L1, A0A0C2LK74, Q99WS0 | 0.653778 | ABC-type transporter, DNA-binding protein, Nucleic acid binding OB-fold tRNA, helicase-type, Antibiotic transport-associated protein, Fatty acid efflux MMPL transporter FarE, RND superfamily resistance-nodulation-cell division:proton (H+) antiporter, 5'-nucleotidase, lipoprotein e(P4) family, Acid phosphatase, Extracellular glutamine-binding protein, Efflux RND transporter permease subunit, MMPL family transporter, RND multidrug efflux transporter Acriflavin resistance protein, Amino acid ABC transporter substrate-binding protein, Cysteine ABC transporter, substrate-binding protein, Transporter substrate-binding domain-containing protein, Amino acid ABC superfamily ATP binding cassette transporter, binding protein, Putative amino acid ABC transporter, Aromatic acid exporter family protein, Membrane protein, ABC transporter permease, Glutamate ABC transporter permease, ABC transporter permease subunit, Arginine-binding protein |
| A0AAJ4YY62 | 0.744 | Gram-positive signal peptide, ysirk family protein |
| A8QKC8 | 0.742 | Neutral metalloproteinase |
| A0A641A8P6 | 0.736 | DNA-binding protein |
| Q9ZFS5, A0A641A9C5, A0AAE8PBR8, Q9RN32, A0A3A5M4N3 | 0.6784 | Exotoxin 1, Superantigen-like protein SSL7, Superantigen-like protein, Exotoxin 11, Superantigen-like protein 7 |
| Q9RL71 | 0.734 | Neutral metalloproteinase |
| A0A0D6GBG7 | 0.728 | FKLRK protein, Gram-positive signal peptide, ysirk family |

| | | |
|---|---|---|
| | | protein, Signal peptide protein |
| A0A0E8IUH3, A0A390UFQ0 | 0.716 | Serine protease |
| Q99TD3, A0A641A7D5, A0AAW4Y9C9 | 0.68 | Haptoglobin receptor A, Iron-regulated surface determinant protein H, Staphylococcus aureus surface protein I |
| A0AAX2K2V4, A0AAW4Y4U5, A0A9P2YZ05 | 0.714 | Surface anchored protein, LPXTG-anchored DUF1542 repeat protein FmtB, DUF1542 domain-containing protein, Methicillin resistance determinant FmtB protein |
| Q6GDD3, A0AAE2ZV88, A0A2U0ITW0, A0A2Y9TLI2 | 0.7095 | Glycerol ester hydrolase 1, Lipase 1, triacylglycerol lipase |
| A0AAJ4YXI7, A0AAE2ZWU, P0C1S5, A0AAE8PAM7, Q6GA99, X5EGV6, Q93PN3, A0AAE8TIW8, W8U414, A0AAE2ZWJ3, A0A2W3XW02, X5DST2, A0AAE8PDN0, A0A166Q5A2, A0A2G4VM73, O87491, A0A641AAJ6, X5DVD1, A0A0G3D1I8, A0A389WRH0, Q6GHV3, A0A0D1GAM5, A0A068DV34, A0A0D6WE06, A0A641A5I4, W8U4R2, A0A641A761, A0A131JAK2, W8U5S1, A0AAE2ZRY0, A0A2C6MZL2, W8URT6, A0A0E1XBG6, Q7BGA5, A0AAE3D5D3, A0AAE8PCK2, | 0.618585 | Iron compound ABC uptake transporter substrate-binding protein, Heme ABC transporter permease, Heme transporter IsdDEF, membrane component IsdD, Iron-regulated surface determinant protein IsdD, Fur-regulated protein A, Iron-regulated surface determinant protein A, Staphylococcal transferrin-binding protein A, High-affinity heme uptake system protein IsdE, Iron-regulated surface determinant protein E, Staphylococcal iron-regulated protein F, Cytochrome aa3-controlling protein, Heme A synthase, ABC transporter substrate-binding protein, Ferric hydroxamate receptor 1, Ferrichrome-binding protein, Iron-hydroxamate ABC transporter substrate-binding protein, Iron(III) dicitrate-binding protein, Iron-dicitrate transporter substrate-binding subunit, FecB, Extracellular solute-binding protein, Periplasmic-iron-binding protein BitC, Iron transporter, Siderophore-mediated iron transport protein, Staphylococcal protein, Iron-regulated surface determinant protein F, Probable heme-iron transport system permease protein IsdF, Staphylococcal iron-regulated protein G, High-affinity iron permease, Iron ($Fe^{3+}$) ABC superfamily ATP binding cassette transporter, binding protein, Iron ABC transporter substrate-binding protein, Iron-siderophore ABC transporter substrate-binding protein, Siderophore staphylobactin ABC transporter, substrate-binding protein SirA, SirA, Staphyloferrin B ABC transporter substrate-binding protein SirA, Lipoprotein, membrane, Putative iron(III) dicitrate transporter binding lipoprotein, Heme transporter IsdA, Iron ($Fe^{2+}$)-regulated surface determinant protein IsdA, Iron-regulated heme-iron binding protein, LPXTG-anchored heme-scavenging protein IsdA, Iron citrate ABC transporter substrate-binding protein, ABC-type siderophore binding protein, Siderophore ABC transporter substrate-binding protein, Ferrichrome ABC transporter subunit, FTR1 family iron permease, Ferric hydroxamate receptor 2, Ferrichrome-binding periplasmic protein, Iron (Fe+3) ABC superfamily ATP binding cassette transporter, Uncharacterized protein sirE, High-affinity $Fe^{2+}$, $Pb^{2+}$ permease-like protein, Heme ABC transporter |

| | | |
|---|---|---|
| A0AAJ4YYM9, Q9X662, Q8KQR2, A0A9P2YW15, A0A3A5LSJ0 | | |
| A0AAX2MHQ7, A0A1S6DAJ0, A0A641A7Q5, A0AAX2MGZ5, A0A9P3DM65, A0AAE8PC44, A0A122JP17, A0AAE8P9X9, X5EF25, Q7WS91, A0AAE8PC88, A0A641A9X6, A0A391DK74, A0A3A3AIJ1, A0AAE8P9T6, A0AAJ4YVK7, A0A0D6HQY2, A0AAE8TKC7, A0A1K9AE69, A0A2W3EU09, A0A0D6GUP8, A0A390ZLD1, A0A641A839, A0A9Q4UZ21, A0A2S6DKY9, A0A0D1HLP4, A0A113XPM5, A0A0D6WBT7, A0AAJ4YVY6, A0A166JQS2, A0A2Y1M5T3, A0AAJ4YYS6, A0A390J584, A0A659I881, W8TVQ4, A0A8G2HYY2, A0A390ZHS0, A0A0D1IQA4, Q2FXH4, A0AAE3D5F7, A0A641A7V7, A0A3M9GHA5, A0AAE8TK32 | 0.63493 | Surface anchored protein, 5'-nucleotidase, LPXTG-anchored adenosine synthase AdsA, LPXTG cell wall anchor domain-containing protein, Cell shape protein MreC, Cell shape-determining protein MreC, Cell wall-anchored protein, Cell-wall-anchored protein SasF, Probable cell wall hydrolase LytN, Cell surface like-protein Map-w, MAP domain-containing protein, Cell-wall-anchored protein SasC, FmtB-like protein, LPXTG-motif cell wall anchor domain-containing protein, Alpha, beta hydrolase, Esterase, lipase, Uncharacterized distant relative of cell wall-associated hydrolases, Adhesin, Truncated MHC class II analog protein, Truncated cell surface protein map-w, Fibronectin-binding protein B, Fibronectin-binding protein FnbB, Cell wall anchored protein, putative, Surface protein, Toxin, Cell wall inhibition responsive protein CwrA, Exported protein, Orthopoxovirus protein, PF05708 family, beta cell surface hydrolase, Extracellular adherence protein Eap, LPXTG cell wall surface anchor family protein, LPXTG-anchored repetitive surface protein SasC |
| P0C1U8, Q6GI34, A0AAE2ZZ37, | 0.6815 | Endoproteinase Glu-C, Glutamyl endopeptidase, Staphylococcal serine proteinase, V8 protease, V8 proteinase, Serine protease |

| | | |
|---|---|---|
| A0A391DGG2 | | |
| A0A641A4T7 | 0.71 | Serine protease |
| A8QKD5 | 0.708 | Neutral metalloproteinase |
| Q9ZFS6 | 0.708 | Exotoxin |
| A0AAX2MJJ6 | 0.708 | Exported protein, Gram-positive signal peptide, ysirk family protein |
| Q53782, A0A163QKA2, A0A0E7RZY7 | 0.68 | Serine protease SplC, Serine protease |
| A0A0D6HFN1, Q53781 | 0.675 | Serine protease, Serine protease SplB |
| A0A0D1JQI9, A0A131K5I5 | 0.698 | Two-component regulator YycH, Two-component system activity regulator YycH, YycH protein |
| A0A0Z0X9F2 | 0.702 | Serine protease |
| A0A641A6X6 | 0.702 | Serine protease |
| A0A0Y9QEI6, P60158, A0A389VIP1, A0A8G2MAE5, A0A2S6D294 | 0.6908 | Immunodominant staphylococcal antigen A, Probable transglycosylase IsaA, SAI-2, Secretory protein SAI-B |
| A0A0D6HF50 | 0.694 | Serine protease |
| A0AAJ4YV38 | 0.692 | Osmotically activated L-carnitine/choline ABC transporter, substrate-binding protein OpuCC |
| A0A641A9M4 | 0.692 | Superantigen-like protein SSL10 |
| A0A5Q2QJY5, Q6G8C4, A0A0D6HF55 | 0.682 | Serine protease, Serine protease SplF |
| A0A0D6W7J6 | 0.69 | Exotoxin, Superantigen-like protein, Superantigen-like protein, exotoxin 14 |
| A0AAN1H0Q7 | 0.69 | Neutral metalloproteinase |
| Q9ZFS3 | 0.688 | Exotoxin 4, Superantigen-like protein, exotoxin 14 |
| A0A6B2ITZ9 | 0.688 | Uncharacterized protein |
| A0A641A901 | 0.686 | FKLRK protein |
| A0AAE2ZQ20 | 0.686 | ABC transporter substrate-binding protein, Periplasmic binding protein |

| | | |
|---|---|---|
| A0AAE8TKE8 | 0.684 | Osmoprotectant ABC transporter substrate-binding protein, Osmotically activated L-carnitine/choline ABC transporter, substrate-binding protein OpuCC |
| A0A389U9K2 | 0.684 | MAP domain-containing protein, Major histocompatibility complex class II analog protein, Map |
| A0A641A6T6 | 0.682 | Osmoprotectant ABC transporter substrate-binding protein, Osmotically activated L-carnitine/choline ABC transporter, substrate-binding protein OpuCC |
| A0A641A9M0, A0A389XVZ5, A0A0D6GAM0 | 0.663333 | Superantigen-like protein SSL5, Superantigen-like protein, Staphylococcal exotoxin |
| Q7X0E4, A0A390PKN6, P0A0L8, P0A0M0, Q8VVS0, Q9F0L7, A0A389VJ56, O33586, A0A2C9TTS8, A0AAE8PC26, A0AAE8PD09, A0A658BFW1, O85383, A0AAE3D531, A0AAE2ZWR5, A0A3A5LV46, P0A0L2, A0A0Z0ILF4, A0A5N5IJZ3, P0A0L5, A0A2P7CQJ7, A0A5S9I2G1, A0A6B3J2A6, A0A6B3INW0, A0A0D3Q467, A0AAE2ZYQ6, A0AAE2ZZJ5, A0A2U0ISP5, A0A4U2KHW9, W8UWT4, Q53643, Q7X0E8 | 0.565875 | Enterotoxin type G, Extracellular enterotoxin type G, Enterotoxin P, phage associated, Staphylococcal enterotoxin type P, SEG, Enterotoxin type H, SEH, Type I toxin-antitoxin system Fst family toxin, Extracellular enterotoxin L, Extracellular enterotoxin type I, Sel, Staphylococcal enterotoxin L, Enterotoxin SEM, Exotoxin, Staphylococcal enterotoxin type I, Staphylococcal enterotoxin type M, Accessory gene regulator D (Pheromone, type II), AgrD, Cyclic lactone autoinducer peptide, Staphylococcal accessory gene regulator protein D, Enterotoxin type I, SEI, Staphylococcal enterotoxin type 26, Type VII secretion system accessory factor EsaA, Enterotoxin type A, SEA, Staphylococcal enterotoxin type O, Enterotoxin type C-3, SEC3, Staphylococcal enterotoxin type 02, Staphylococcal enterotoxin type K, Staphylococcal enterotoxin type Q, Enterotoxin, Staphylococcal enterotoxin type C1, Staphylococcal enterotoxin type U, SEN, Staphylococcal enterotoxin type N, Autoinducer peptide, Pre-pheromone, Regulator protein D, Regulator protein agrD, Enterotoxin SEI variant |
| A0A389VJ67 | 0.678 | Beta-lactamase, DUF4888 domain-containing protein |
| A0A391DDB2 | 0.676 | Glycine betaine/L-proline ABC superfamily ATP binding cassette transporter, binding protein, Osmoprotectant ABC transporter substrate-binding protein, Osmotically activated L-carnitine/choline ABC transporter, substrate-binding protein OpuCC |

| | | |
|---|---|---|
| Q9ZFS4 | 0.676 | Exotoxin 5 |
| A0A3A3AEZ5 | 0.672 | Exported protein, FKLRK protein |
| A0A641A9T1 | 0.67 | Serine protease |
| Q6GGX3 | 0.67 | ECM-binding protein homolog, Extracellular matrix-binding protein ebh |
| Q2FYJ6 | 0.67 | ECM-binding protein homolog, Extracellular matrix-binding protein ebh |
| A0A0D6GGQ7 | 0.668 | Peptide ABC transporter substrate-binding protein |
| A0A131K2A0 | 0.664 | Lipoprotein, Polysaccharide deacetylase family protein |
| A0A0D6HBZ8, A0A641A4L8, A0A2S6DNW9, A0A390EPY9 | 0.6555 | CamS family sex pheromone protein, Lipoprotein, Lipoprotein (Pheromone), Pheromone lipoprotein CamS |
| A0A390B0Z6 | 0.664 | Membrane lipoprotein, Staphylococcal lipoprotein, Tandem-type lipoprotein |
| A0A2C9TU38 | 0.662 | MAP domain-containing protein, Protein map |
| A0A641A5J8 | 0.662 | Serine protease |
| X5ELQ6 | 0.66 | Lipoprotein, Polysaccharide deacetylase |
| A0AAJ4YZ65 | 0.66 | MHC class II antigen-like protein |
| A0AAE8PBN0 | 0.658 | Superantigen-like protein SSL10, Superantigen-like protein, exotoxin 14 |
| Q9EY53 | 0.658 | RGD-containing lipoprotein |
| A0A2S6D1X8 | 0.658 | Osmoprotectant ABC transporter substrate-binding protein, Osmotically activated L-carnitine/choline ABC transporter, substrate-binding protein OpuCC |
| A0AAJ5CTI3 | 0.656 | Superantigen-like protein |
| A0AAE3D6E5 | 0.654 | Lipoprotein, Polysaccharide deacetylase |
| A0AAJ4YZ59 | 0.654 | Enterotoxin P, phage associated, Exotoxin |
| A0AAE8TJ68 | 0.654 | Peptide ABC transporter substrate-binding protein, RGD-containing lipoprotein |
| A0AAJ4YVY9 | 0.654 | Serine protease |
| A0A0D3QAE7 | 0.654 | Glycine/betaine ABC transporter substrate-binding protein, Osmoprotectant ABC transporter substrate-binding protein, Osmotically activated L-carnitine/choline ABC transporter, substrate-binding protein OpuCC |

| | | |
|---|---|---|
| A0A0Z4ZGY7, Q79SZ7, A0AAE8TIS4, P10335 | 0.649 | triacylglycerol lipase, Glycerol ester hydrolase 2, Lipase 2 |
| Q2G1F3 | 0.652 | Uncharacterized protein SAOUHSC_00172 |
| A0AAE2ZTC3 | 0.652 | Membrane lipoprotein, Tandem-type lipoprotein |
| A0A9P2YWD9 | 0.652 | C4-dicarboxylate anaerobic carrier |
| Q6GJX5, A0A641A8S4, A0A0B4N958 | 0.638667 | Efem, EfeO family lipoprotein |
| A0A641AA65 | 0.65 | triacylglycerol lipase |
| A0AAJ4YY44 | 0.65 | Lipoprotein (Pheromone) |
| A0A389PUB8 | 0.65 | ABC-type dipeptide transport system periplasmic component-like protein, Peptide ABC transporter substrate-binding protein, RGD-containing lipoprotein |
| A0A0D6HDR6 | 0.65 | Serine protease |
| A0A0D6HDD7 | 0.648 | Beta-lactamase, DUF4888 domain-containing protein |
| A0A2I7YBL5, H9BJ68, H9BJ77 | 0.626667 | YfcC family protein, ArcD-like protein, Arginine, ornithine antiporter ArcD |
| A0A641A9B4 | 0.644 | Lipoprotein, Polysaccharide deacetylase |
| A0AAJ5CT13 | 0.644 | Ferrichrome-binding periplasmic protein |
| Q2G2B2 | 0.642 | Surface protein G |
| A0A641AC18 | 0.642 | FPRL1 inhibitory protein |
| A0A6B2IY43 | 0.64 | Exotoxin |
| Q6YCN3 | 0.638 | Enterotoxin SeN |
| A0A166EAB0 | 0.638 | Lipoprotein |
| A0AAX2K114 | 0.634 | Exotoxin 13 |
| A0A641ABQ3 | 0.634 | DUF1672 domain-containing protein |
| Q6YCN4 | 0.634 | Enterotoxin B |
| A0A166DEL7 | 0.634 | Exported protein, Secretory extracellular matrix and plasma binding protein |
| A0A166JSW2 | 0.632 | Exported protein |

| | | |
|---|---|---|
| A0AAJ4YXL0 | 0.63 | Bifunctional autolysin |
| A0AAE2ZW53 | 0.63 | Staphylococcal exotoxin, Superantigen-like protein |
| A0A0D3Q6D3 | 0.63 | FPRL1 inhibitory protein |
| A0AAJ4YYA4, W8TMJ4, A0A380E1J8 | 0.589333 | Iron-regulated surface determinant protein C, Staphylococcal iron-regulated protein D |
| A0A390ZCI1 | 0.628 | DUF1672 domain-containing protein, DUF1672 family protein, Lipoprotein |
| A0A2U0IR43 | 0.628 | FPRL1 inhibitory protein |
| A0A3A3ALE1 | 0.628 | Bifunctional autolysin |
| Q33C62 | 0.628 | Lipoprotein, Similar to LMP group of surface-lacated membrane protein |
| H9BJ80 | 0.626 | ArcD-like protein |
| A0A4T9YI84 | 0.626 | Staphylococcal complement inhibitor |
| A0AAJ4YZ26, A0AAE8PAT4, A0A0D1GC10 | 0.612667 | NERD domain-containing protein |
| Q6GAG0 | 0.624 | Bifunctional autolysin |
| A0AAE2ZZL2 | 0.622 | Extracellular adherence protein Eap/Map, MAP domain-containing protein |
| A0AAE2ZW11 | 0.622 | Exotoxin, Superantigen-like protein |
| Q2FZK7 | 0.622 | Bifunctional autolysin |
| A0A641AAF9, A0A390F740 | 0.617 | Superantigen-like protein SSL9, Exotoxin |
| A0A0D3Q4X2, A0A2W3PXW4 | 0.6 | Lipoprotein |
| A0A0D1H8K9 | 0.62 | Exported protein, Peroxidase inhibitor |
| A0AAE8PCR4 | 0.618 | Bifunctional autolysin |
| A0A0D6G9V6 | 0.618 | Exotoxin, Superantigen-like protein |
| A0AAX2JZY5 | 0.616 | Collagen adhesin |
| A0AAE2ZTI9 | 0.616 | Staphylococcal complement inhibitor |
| A0A3A3AD99, A0A6A9GVV2, X5DU72 | 0.602 | Phosphate, phosphite, phosphonate ABC transporter substrate-binding protein, phosphonate ABC transporter periplasmic protein, Phosphonate ABC transporter |

| | | |
|---|---|---|
| | | phosphate-binding periplasmic component, phosphonate ABC transporter, periplasmic binding family protein, Phosphonate ABC transporter substrate-binding protein |
| A0A0D6GBL1 | 0.616 | Exotoxin, Superantigen-like protein |
| A0AAE8PAL3 | 0.614 | Lipoprotein |
| O54462 | 0.614 | Superantigen-like protein, exotoxin 14, Toxic shock syndrome toxin TSST-1, Toxic shock syndrome toxin-1 |
| A0A391M7N5, A0AAE8TK88 | 0.604 | Cell-wall binding lipoprotein, YkyA, Lipoprotein, YkyA family protein |
| A0AAJ4YYP0 | 0.614 | Lipoprotein |
| A5JNM2 | 0.61 | Enterotoxin M, Enterotoxin SEM |
| A0A0D6WG27 | 0.61 | Staphylococcal complement inhibitor |
| A0A641AAE0 | 0.61 | Uncharacterized protein |
| X5E0C0 | 0.608 | Lipoprotein |
| A0AAJ4YYD7 | 0.608 | Lipoprotein |
| A0A0D6HYB8 | 0.606 | Extracellular ECM and plasma binding protein Emp, Extracellular matrix protein-binding adhesin Emp, Extracellular matrix protein-binding protein emp |
| A0A7U7IED2 | 0.606 | Collagen adhesin |
| A0AAE2ZTW7 | 0.606 | DUF1672 domain-containing protein |
| A0A3A2ILB4 | 0.606 | Extracellular ECM and plasma binding protein Emp, Extracellular matrix protein-binding adhesin Emp, Secretory extracellular matrix and plasma binding protein |
| A0AAE2ZWL4 | 0.604 | FPRL1 inhibitory protein |
| A0A641A8E7 | 0.602 | Membrane lipoprotein, Tandem-type lipoprotein |
| A0AAJ5CRW3, A0A2C9TY49 | 0.58 | Lipoprotein, DUF1672 domain-containing protein, DUF1672 family protein |
| A0A641A6V2 | 0.602 | Extracellular matrix protein-binding adhesin Emp |
| A0A0H2DUF0 | 0.6 | Staphylococcal complement inhibitor |
| A0A0D6HEH5 | 0.6 | DUF445 domain-containing protein, DUF445 family protein, Membrane associated protein, Membrane protein |
| Q83TG5, A0A2L2E2K2, A0A9Q3MR89 | 0.595333 | DM13 domain-containing protein, Lipoprotein, SAR0761-like protein |

| | | |
|---|---|---|
| Q9LAB5, A0AAE2ZY64, A0A162H218 | 0.598 | Immunodominant staphylococcal antigen B |
| Q6GED5 | 0.598 | Staphylococcal secretory antigen ssaA2 |
| A0A166K1P5, A0A2U0ITH7, A0A0D6GPA6, A0A641A876, A0AAE3D5U9 | 0.5844 | Accessory Sec system protein Asp3, Accessory secretory protein Asp3 |
| A0A166EPD7 | 0.594 | Staphylococcal complement inhibitor |
| A0AAE8TIB6 | 0.594 | Extracellular ECM and plasma binding protein Emp, Extracellular matrix protein-binding adhesin Emp |
| A0A6B5KXH1 | 0.594 | Lipoprotein |
| X5EHQ5 | 0.594 | Uncharacterized protein |
| A0A9Q7J6Z3 | 0.592 | DUF445 domain-containing protein, DUF445 family protein, Membrane protein |
| A0A162FG49 | 0.592 | Lipoprotein |
| A0A389QEW1 | 0.592 | DUF1672 domain-containing protein, DUF1672 family protein, Lipoprotein |
| A0A390BBS7 | 0.592 | Lipoprotein |
| A0AAE3D601 | 0.588 | DUF1672 domain-containing protein, Lipoprotein |
| X5DRS2 | 0.588 | Lipoprotein |
| A0AAE3D3F5 | 0.588 | Lipoprotein |
| A0A641A9L3 | 0.588 | Superantigen-like protein SSL1 |
| A0AAE8TJG1 | 0.586 | Genomic island nu Sa alpha2 |
| W8U6M5 | 0.586 | Lipoprotein |
| A0A390ZN89 | 0.586 | Exotoxin, Superantigen-like protein |
| A0A068DWK6 | 0.586 | DUF4889 domain-containing protein |
| A0A0D6I082 | 0.584 | Extracellular matrix and plasma binding protein, von Willebrand binding protein |
| A0A2I7YAS3, A0A659ID04 | 0.565 | DMT family transporter, EamA family transporter |
| A0A227LZW5 | 0.58 | Lipoprotein |
| A0AAE8TIW2 | 0.578 | Myeloperoxidase inhibitor SPIN |

| | | |
|---|---|---|
| A0AAJ4YX88 | 0.578 | Membrane associated protein |
| A0A131HYC7 | 0.574 | Peptidase, Subtilase family protease |
| A0A2L2E3D0 | 0.572 | Exported protein |
| A0A641ABK1 | 0.57 | DUF1672 domain-containing protein |
| A0A641A6G6 | 0.57 | DUF445 family protein |
| W8TRN2 | 0.568 | Exported protein |
| A0AAE3D6H3 | 0.566 | DUF445 domain-containing protein, DUF445 family protein, Membrane associated protein, Membrane protein |
| A0A9P2YV90 | 0.562 | Lipoprotein |
| A0A659ZYI3 | 0.562 | Lipoprotein |
| Q2FVL6 | 0.562 | DUF4467 domain-containing protein |
| A0AAE2ZZC2 | 0.558 | DUF1672 domain-containing protein |
| W8U0W4 | 0.554 | Lipoprotein |
| A0A131JS96 | 0.554 | Exported protein |
| A0A0D6HLH7 | 0.554 | DUF1672 domain-containing protein, DUF1672 family protein, Lipoprotein, Putative lipoprotein with DUF1672 |
| A0A0D1GQ07 | 0.554 | Exported protein |
| A0A641ACA7 | 0.55 | Lactococcin 972 family bacteriocin |
| A0A391E4G2 | 0.544 | DUF4889 domain-containing protein |
| A0AAE2ZXF7 | 0.534 | DUF4889 domain-containing protein |
| Q7DI43 | 0.53 | Chitinase, DUF5011 domain-containing protein, Ig-like domain (Group 3) protein, IraE protein, Putative BIG_3-domain protein |
| A0A0G2LTE5 | 0.522 | Lantibiotic |
| A0A0D3Q8P9 | 0.522 | Lantibiotic |
| A0A163VMG3 | 0.52 | Lipoprotein |
| A0AAW4Y5F4 | 0.516 | Exported protein |
| Q99RL9 | 0.516 | DUF4467 domain-containing protein, Lipoprotein, SA2198 |
| A0A166J6C2 | 0.512 | Pathogenicity island protein |
| X5DWZ3 | 0.51 | FeoB-associated Cys-rich membrane protein, Virus attachment p12 family protein |
| A0A0D1IZ30 | 0.51 | DUF1648 domain-containing protein, Predicted membrane |

| | | |
|---|---|---|
| | | protein |
| H9BRQ6 | 0.51 | Phenol-soluble modulin PSM-alpha-2, Psm alpha-2 |
| A0AAE8PD46 | 0.51 | DUF1648 domain-containing protein, Predicted membrane protein |
| A0AAJ5CRM5 | 0.508 | Lipoprotein |
| A0A0D6GLP7 | 0.504 | DUF1648 domain-containing protein |
| A0A9P2YYE8 | 0.504 | Uncharacterized protein |
| A0AAJ4YZG0 | 0.502 | Conserved low-complexity protein |
| A0A266CVG1 | 0.5 | Uncharacterized protein |
| A0AAE2ZV66 | 0.496 | Low-complexity protein |
| A0A162H7Z8 | 0.49 | FeoB-associated Cys-rich membrane protein, Virus attachment p12 family protein |
| B0I1V8 | 0.488 | Bacteriocin, lactococcin 972 family protein, Lactococcin 972 family bacteriocin, Lactococcin-related protein, Putative bacteriocin |
| A0A2C9TMN6 | 0.486 | Exported protein |
| A0A0D1IH94 | 0.486 | Membrane protein |
| A0A3A3AKD2 | 0.486 | FeoB-associated Cys-rich membrane protein, Virus attachment p12 family protein |
| H9BRQ5 | 0.486 | Phenol-soluble modulin PSM-alpha-1, Phenol-soluble modulin alpha 1 peptide, Psm alpha-1 |
| Q1XG25 | 0.484 | Uncharacterized protein |
| A0A0D3QAT6 | 0.482 | Membrane protein |
| A0A3A1VUR5 | 0.482 | DUF1648 domain-containing protein, Predicted membrane protein |
| A0AAE2ZSI2 | 0.48 | Membrane protein |
| A0AAE2ZU24 | 0.478 | Serine protease |
| A0A389UAT8 | 0.464 | Uncharacterized protein |
| A0A0D6HYP4 | 0.392 | Uncharacterized protein |
| D5MS78 | 0.372 | Phenol-soluble modulin PSM-mec, Phenol-soluble modulin-mec, Psm-mec protein |

**Supplementary Table 10: Antigen predicted *S. aureus* proteins that were removed due to high human protein homology (BLASTp percent identity > 30% (e-value < 0.05) or BLASTp e-value < $10^{-6}$).** A total of 24 proteins.

| Accession(s) | Antigen Probability | Protein Names |
|---|---|---|
| A0A390ZED4, H6UH57, A0AAX2K025 | 0.725 | Alkaline phosphatase |
| A0A6B5FIU0 | 0.732 | Alkaline phosphatase |
| W8U0B2, A0A167H2Q7 | 0.697 | Glycerophosphodiester phosphodiesterase, Glycerophosphoryl diester phosphodiesterase |
| A0AAJ4YVQ8, A0A090LRP9, W8TV96 | 0.649 | Foldase protein PrsA |
| A0AAE2ZZ06 | 0.666 | Beta-lactamase family protein, Flp protein, Serine hydrolase FLP |
| A0AAJ4YVE5 | 0.662 | Beta-lactamase family protein |
| A0A0D6HEZ2 | 0.654 | Leader peptide-processing serine protease |
| Q6GDH2, Q2FUY2, A0A2U0ITG8, Q6G644 | 0.614 | Clumping factor B, Fibrinogen receptor B, Fibrinogen-binding protein B, MSCRAMM family adhesin clumping factor ClfB |
| A0AAJ5CTN7 | 0.65 | 5'-nucleotidase |
| A0A166LM76 | 0.646 | Cysteine ABC transporter, substrate-binding protein |
| Q9KJ74 | 0.626 | FmtA-like protein, Protein flp |
| A0A641A8R9 | 0.62 | Protein flp |
| Q99RD2, Q6GDU5, A0A7K3N1M9, A0AAX2K2C1 | 0.614 | Fibronectin-binding protein A, Fibronectin-binding protein FnbA, Fibronectin binding protein FnbA |
| A0AAE8P9K4 | 0.602 | Cysteine ABC transporter, substrate-binding protein, Transporter substrate-binding domain-containing protein |
| A0AAE8TK43, A0A0D6GUT9, A0AAJ4YWR0 | 0.591 | 4, 4'-diapophytoene desaturase (4, 4'-diaponeurosporene-forming), Dehydrosqualene desaturase |
| A0A0D3Q629, W8U0F6 | 0.581 | DUF21 domain-containing protein, Hemolysin, HlyC, CorC family transporter, CNNM domain-containing protein, Mg2+ and Co2+ transporter, CorB |
| D2J6U4 | 0.588 | Multi-copper oxidase Mco, Multicopper oxidase |
| A0AAE2ZW9, | 0.548 | 4, 4'-diaponeurosporenoate glycosyltransferase |

| | | |
|---|---|---|
| A0A391DD44, A0A0Y9ABM, A0A0D6GUE2 | | |
| A0AAJ5CS60 | 0.586 | Esterase/lipase |
| A0A641A9P8 | 0.578 | 4,4'-diapophytoene desaturase (4,4'-diaponeurosporene-forming), Dehydrosqualene desaturase |
| W8TMQ6 | 0.574 | Probable quinol oxidase subunit 2, Quinol oxidase polypeptide II |
| W8UBK2 | 0.558 | succinate dehydrogenase |
| A0AAX2ML76 | 0.498 | UDP-N-acetylglucosamine--N-acetylmuramyl-(pentapeptide) pyrophosphoryl-undecaprenol N-acetylglucosamine transferase, Undecaprenyl-PP-MurNAc-pentapeptide-UDPGlcNAc GlcNAc transferase |
| W8USZ3 | 0.336 | DUF423 domain-containing protein, Membrane protein, Membrane spanning protein |

**Supplementary Table 11: Protein structure availability in category compared to number of antigens and non-antigen random proteins.** PDB files are sourced from the Protein Data Bank (PDB) and AlphaFold Protein Structure Database.

| Bacterium | No. of PDB files for Antigens | No. of Antigens | No. of PDB files for Random Proteins | No. of Random Proteins |
|---|---|---|---|---|
| B.melitensis | 107 | 138 | 133 | 200 |
| C.burnetii | 141 | 148 | 202 | 200 |
| C.trachomatis | 29 | 123 | 1 | 200 |
| H.influenzae | 80 | 77 | 19 | 200 |
| H.pylori | 48 | 32 | 10 | 200 |
| N.gonorrhoeae | 100 | 87 | 211 | 200 |
| P.aeruginosa | 59 | 129 | 4 | 200 |
| S.enteritidis | 27 | 28 | 4 | 200 |
| S.pneumoniae | 51 | 69 | 0 | 200 |
| S.pyogenes | 48 | 132 | 3 | 197 |
| T.pallidum | 140 | 136 | 166 | 200 |
| V.cholerae | 157 | 140 | 204 | 200 |
| Total | 987 | 1239 | 957 | 2397 |

32. Beare PA, Unsworth N, Andoh M, Voth DE, Omsland A, Gilk SD, et al. Comparative genomics

reveal extensive transposon-mediated genomic plasticity and diversity among potential effector proteins within the genus Coxiella. Infect Immun. 2009 Feb;77(2):642–56.

49. Nezafat N, Karimi Z, Eslami M, Mohkam M, Zandian S, Ghasemi Y. Designing an efficient

multi-epitope peptide vaccine against Vibrio cholerae via combined immunoinformatics and protein interaction based approaches. Comput Biol Chem. 2016 Jun;62:82–95.

65. Hallgren J, Tsirigos KD, Pedersen MD, Almagro Armenteros JJ, Marcatili P, Nielsen H, et al. DeepTMHMM predicts alpha and beta transmembrane proteins using deep neural networks

[Internet]. bioRxiv. 2022 [cited 2025 Nov 19]. p. 2022.04.08.487609. Available from: https://www.biorxiv.org/content/10.1101/2022.04.08.487609v1.abstract
